# Distinct guard cell specific remodeling of chromatin accessibility during abscisic acid and CO2 dependent stomatal regulation

**DOI:** 10.1101/2023.05.11.540345

**Authors:** Charles A. Seller, Julian I. Schroeder

## Abstract

In plants, epidermal guard cells integrate and respond to numerous environmental signals to control stomatal pore apertures thereby regulating gas exchange. Chromatin structure controls transcription factor access to the genome, but whether large-scale chromatin remodeling occurs in guard cells during stomatal movements, and in response to the hormone abscisic acid (ABA) in general, remain unknown. Here we isolate guard cell nuclei from *Arabidopsis thaliana* plants to examine whether the physiological signals, ABA and CO_2_, regulate guard cell chromatin during stomatal movements. Our cell type specific analyses uncover patterns of chromatin accessibility specific to guard cells and define novel cis-regulatory sequences supporting guard cell specific gene expression. We find that ABA triggers extensive and dynamic chromatin remodeling in guard cells, roots, and mesophyll cells with clear patterns of cell-type specificity. DNA motif analyses uncover binding sites for distinct transcription factors enriched in ABA-induced and ABA-repressed chromatin. We identify the ABF/AREB bZIP-type transcription factors that are required for ABA-triggered chromatin opening in guard cells and implicate the inhibition of a set of bHLH-type transcription factors in controlling ABA-repressed chromatin. Moreover, we demonstrate that ABA and CO_2_ induce distinct programs of chromatin remodeling. We provide insight into the control of guard cell chromatin dynamics and propose that ABA-induced chromatin remodeling primes the genome for abiotic stress resistance.

**Significance statement:** Specialized leaf cells called guard cells integrate environmental cues to optimally control the size of microscopic stomatal pores. The hormone abscisic acid (ABA), a key regulator of plant drought responses, and changes in atmospheric CO_2_ concentration are signals that control stomatal aperture size, but whether these signals also regulate genome packaging into chromatin is unknown. Using guard cell specific chromatin profiling we uncovered regulatory DNA sequences driving specific gene expression in this cell-type. We also discovered that ABA triggers extensive and persistent changes to chromatin structure in guard cells. Unexpectedly, exposure of plants to elevated atmospheric CO_2_ had only minimal impact on chromatin dynamics. Furthermore, we identified the specific transcription factors that regulate ABA-induced chromatin dynamics in guard cells.

## INTRODUCTION

Organisms evolved mechanisms that connect the activity of their genome to the conditions in their environment. Developmental and environmental signals can impact the genome by regulating transcription factor (TF) binding to cognate DNA sequences known as cis-regulatory elements (CRE) (1). In eukaryotes, this regulation occurs in the context of chromatin where nucleosomes can impede transcription factor binding to target sequences (2, 3). Consequently, the remodeling of chromatin structure to allow access of transcription factors to target DNA is thought to be a key step in gene regulation. Plant hormones are key signaling molecules controlling numerous aspects of plant life that coordinate genome activity with environmental conditions (4, 5). Recent studies have provided insight into how chromatin structure changes during plant development (6–12) and in response to environmental and hormonal stimuli (13–15).

Here we focus on the abscisic acid (ABA) signal transduction pathway to explore the connection between hormone signaling and chromatin structure in plants. ABA is a major plant stress hormone that accumulates in cells and tissues experiencing abiotic stress, especially those linked to plant water-status (16–18). ABA signals through a well understood core module consisting of PYR/PYL/RCAR ABA receptor proteins, PP2C protein phosphatases, and SnRK2 protein kinases (19–21). Upon ABA binding, PYR/PYL/RCAR receptor proteins inactivate PP2C phosphatases resulting in the derepression of SnRK2 kinases. ABA-activated SnRK2 kinases then phosphorylate and thereby directly regulate downstream proteins including numerous transcription factors (22–28). Consequently, ABA triggers the differential expression of thousands of genes in the *Arabidopsis thaliana* genome (4, 29–31). However, it remains unclear, especially at the genome scale, whether ABA signaling reshapes chromatin structure, and how ABA-regulated TFs might function in the context of chromatin.

One well known action of ABA is to trigger the closure of the stomatal aperture (17). Stomata are small pores on the surface of the leaf that mediate gas exchange and are formed by a pair of specialized epidermal cells known as guard cells. Guard cells perceive and respond to multiple environmental and hormonal cues in order to optimally regulate the size of the stomatal pore (32–34). For instance, some signals such as elevated atmospheric carbon dioxide (CO_2_), ABA, and the immune elicitor Flg22 trigger stomatal closure, while reduced CO_2_, light, and heat trigger stomatal opening. Because of their exquisite environmental reactivity, guard cells are an excellent model cell system to investigate how different physiological signals regulate chromatin structure in a cell-type specific manner. Although the recent application of single-cell RNA-seq to *Arabidopsis* has expanded our understanding of gene regulation, these studies often focus on plate-grown seedlings (35, 36) and/or resulted in datasets with limited representation of guard cells (37, 38). To apply epigenomic techniques to the question of how guard cells respond to different stimuli new protocols are needed that can isolate large numbers of pure guard cells from soil grown plants, a more relevant setting for physiological studies.

In this paper, we describe the development of an approach to purify guard cell nuclei and by deploying this method, uncover how guard cell chromatin structure changes in response to different stimuli that drive stomatal movements. We profiled the chromatin and transcriptional reprogramming in response to ABA in three different developmental contexts – roots, mesophyll cells, and guard cells. We map thousands of loci that gain or lose chromatin accessibility in response to ABA and link these regions to co-regulated transcripts, uncovering a critical role for chromatin dynamics in controlling ABA-dependent transcription. Furthermore, we show that genome-wide and persistent changes to chromatin structure accompany ABA-induced stomatal closure and that four related bZIP transcription factors known as ABRE Binding Factors (ABFs) are required for initiating the majority of ABA-induced chromatin opening. In contrast, we implicate a family of related bHLH-type TFs known as ABA-Kinase Substrates (AKSs) in the maintenance of open chromatin upstream of ABA-repressed genes. Finally, we show that in guard cells, changes to atmospheric CO_2_-concentration trigger distinct and more limited chromatin remodeling and gene regulatory programs from those mediated by abscisic acid.

## RESULTS

### ABA induces rapid and genome-wide chromatin remodeling in roots

As an initial test of the relationship between ABA and chromatin we focused on *Arabidopsis thaliana* seedlings. To measure changes to chromatin structure we combined fluorescence activated nuclei sorting (FANS) with the Assay for Transposase-Accessible Chromatin with sequencing (ATAC-seq), a quantitative measurement of chromatin accessibility across the genome (39, 40). 50,000 nuclei per sample were isolated using FANS (Fig. S1A). We called 25,753 ATAC-seq peaks across all whole seedling and seedling root libraries (Datasets S1 and S2). Genomic regions coinciding with these peaks are known as Accessible Chromatin Regions (ACRs) and predominantly aligned with upstream regulatory regions in the genome (Fig. S1B-E). In contrast, ATAC-seq reads derived from purified genomic DNA did not cluster into defined peaks (Fig. S1B,C). We next treated whole seedlings with ABA and profiled chromatin accessibility after 4 hours. Differential analysis revealed hundreds of regions that significantly gain and lose accessibility in response to ABA (Fig. 1A, Dataset S3).

**Figure 1.**
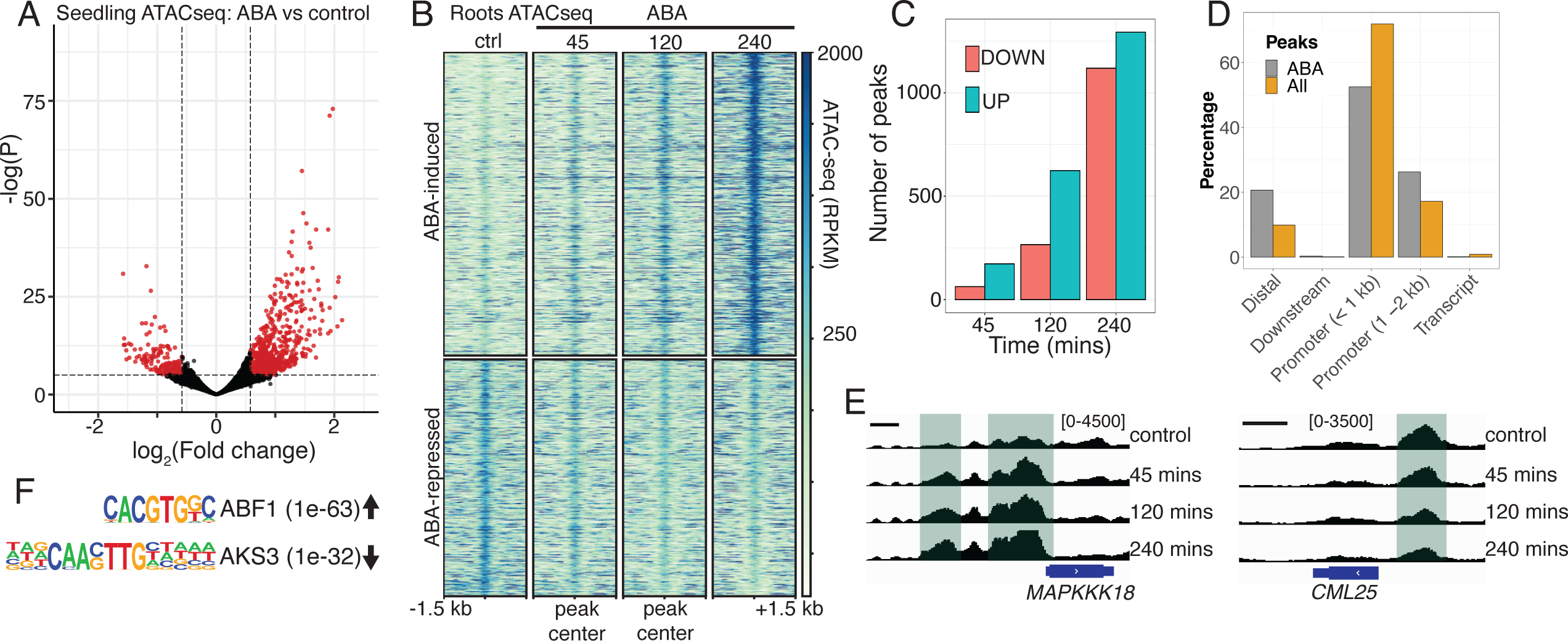
ABA induces rapid and genome-wide chromatin remodeling in seedlings. **A)** Volcano plot of differential ATAC-seq analysis showing regions that significantly change in chromatin accessibility in seedlings after 4hrs of ABA treatment. ABA increased accessibility at 666 regions and decreased accessibility at 214 regions (FDR < 0.001 and FC > 1.5, differential peaks colored in red). **B)** Heatmap of root nuclei ATAC-seq signal (RPKM-normalized) centered over ATAC-seq peaks (+/- 1.5 kb) with ABA-regulated chromatin accessibility during the indicated ABA treatment time course. Across all timepoints, ABA increased accessibility at 1293 regions and decreased accessibility at 1119 regions (FDR < 0.001 and FC > 1.5). **C)** Number of ABA-regulated ATAC-seq peaks over time in roots. ABA-induced peaks are colored in blue while ABA-repressed peaks are colored in pink. **D)** Distribution among annotated genomic features of either all ATAC-seq peaks (orange) or ABA-regulated ATAC-seq peaks (grey). **E)** Genome browser snapshots at two representative genes (*MAPKKK18* and *CML25*) showing changes in ATAC-seq signal in upstream regions following ABA treatment. Scale bars indicate 500 bp. **F)** The transcription factor binding motifs with the highest enrichment (by binomial p-value) in ABA-induced and ABA-repressed chromatin regions in roots.

To capture the dynamics of ABA-induced chromatin remodeling we focused on roots which can rapidly take up ABA from their surroundings (41, 42). Using FANS, we isolated root nuclei at three different timepoints following ABA treatment (after 45, 120, and 240 minutes) and performed ATAC-seq. Differential analysis revealed that ABA triggers time-dependent changes in chromatin accessibility in roots (Dataset S3). Heatmaps centered over these ABA regulated ACRs illustrate the progressive impact of ABA on chromatin structure, with more regions of the genome changing in ATAC-signal over time (Fig. 1B,C). We observed significant changes to chromatin after only 45 minutes of ABA treatment, with 163 ACRs located in upstream regulatory regions gaining accessibility and 64 losing accessibility (Fig. 1C,E). After 4 hours, ABA had activated 1293 ACRs and repressed 1119 ACRs (Dataset S3). Interestingly, we found that ABA-regulated ACRs tend to be located further away from transcription start sites (> 1kb) than static regions (Fig. 1D). ABA-induced ACRs were associated with downstream genes enriched for gene ontology (GO) terms such as “response to water deprivation”, “response to osmotic stress”, and “response to abscisic acid” (Dataset S6).

To explore the role of specific transcription factors in orchestrating these changes we performed DNA motif analysis using a set of motifs empirically generated by the in vitro DNA binding assay DAP-seq (43). We found that motifs recognized by ABF/AREB transcription factors such as ABF1/2/3/4 and ABI5 were highly enriched in ABA-induced ACRs (Fig. 1F, example ABF1, p = 1e-63, Dataset S7). In contrast, we found that ABA-repressed ACRs showed enrichment for motifs recognized by the related bHLH type transcription factors AKS1, AKS3, and bHLH80 (Fig. 1F, example AKS3, p = 1e-32, Dataset S7). Notably, both ABF1/2/3/4 and AKS1/2/3 proteins are direct targets of ABA-regulated SnRK2 kinases (22, 24). Previous research has shown a role for ABA-dependent inhibition of bHLH-type AKS transcription factors in the transcriptional repression of the guard cell K^+^ channel *KAT1* (24, 44). Interestingly, our findings suggest a larger role for the AKS clade of closely related bHLH proteins in the regulation of ABA-repressed genes. We conclude that ABA can trigger rapid and genome-wide chromatin remodeling in root tissue.

### Revealing guard cell specific patterns of chromatin accessibility

Encouraged by our findings in whole seedlings and roots we next decided to investigate the effect of ABA on chromatin in mature guard cells. We first developed a FACS-based strategy to purify guard cell nuclei from soil-grown plants. Nuclei were labeled by expressing the histone H2B fused to GFP under the control of the guard cell *pGC1* promoter (45). We first confirmed the guard cell specificity of the H2B-GFP fusion in our transgenic lines using fluorescence microscopy (Fig. 2A) and confirmed that ABA treatment did not alter the guard cell specific expression (Fig. S2A). We first isolated nuclei from leaf epidermal samples harvested from *GC1:H2B-GFP* plants. DAPI stained nuclei preparations were then analyzed using FACS (Fig S2B). We could define a clearly separated GFP-positive population among all detected nuclei (Fig. 2B, Fig. S2C). In contrast to many other cell types in the *Arabidopsis* leaf, guard cells are diploid, and GFP-positive nuclei from *pGC1:H2B-GFP* expressing plants had a 2n DNA content while GFP-negative nuclei were distributed among higher ploidy levels (Fig. 2C). Approximately 5% of GFP-positive particles (41 out of 5037 total from a single FACS run) had DAPI content greater than 2N, however we sorted only 2N nuclei for downstream applications. This method allowed us to obtain ∼30,000 guard cell nuclei with an estimated purity of 98% (Fig. S2D) from the leaves of 40-50 plants. To assay guard cell chromatin structure, nuclei isolated using this method were used to generate ATAC-seq libraries. The resulting libraries were of high quality with clear concentration of peaks in annotated upstream regulatory regions (Fig. S2E), enrichment of open chromatin fragments (< 100 bp) immediately upstream of transcription start sites (Fig. S2F), strong correlation between biological replicates (Fig. S2G), and high FRiP (fraction of reads in peaks) scores (Dataset S1).

**Fig 2.**
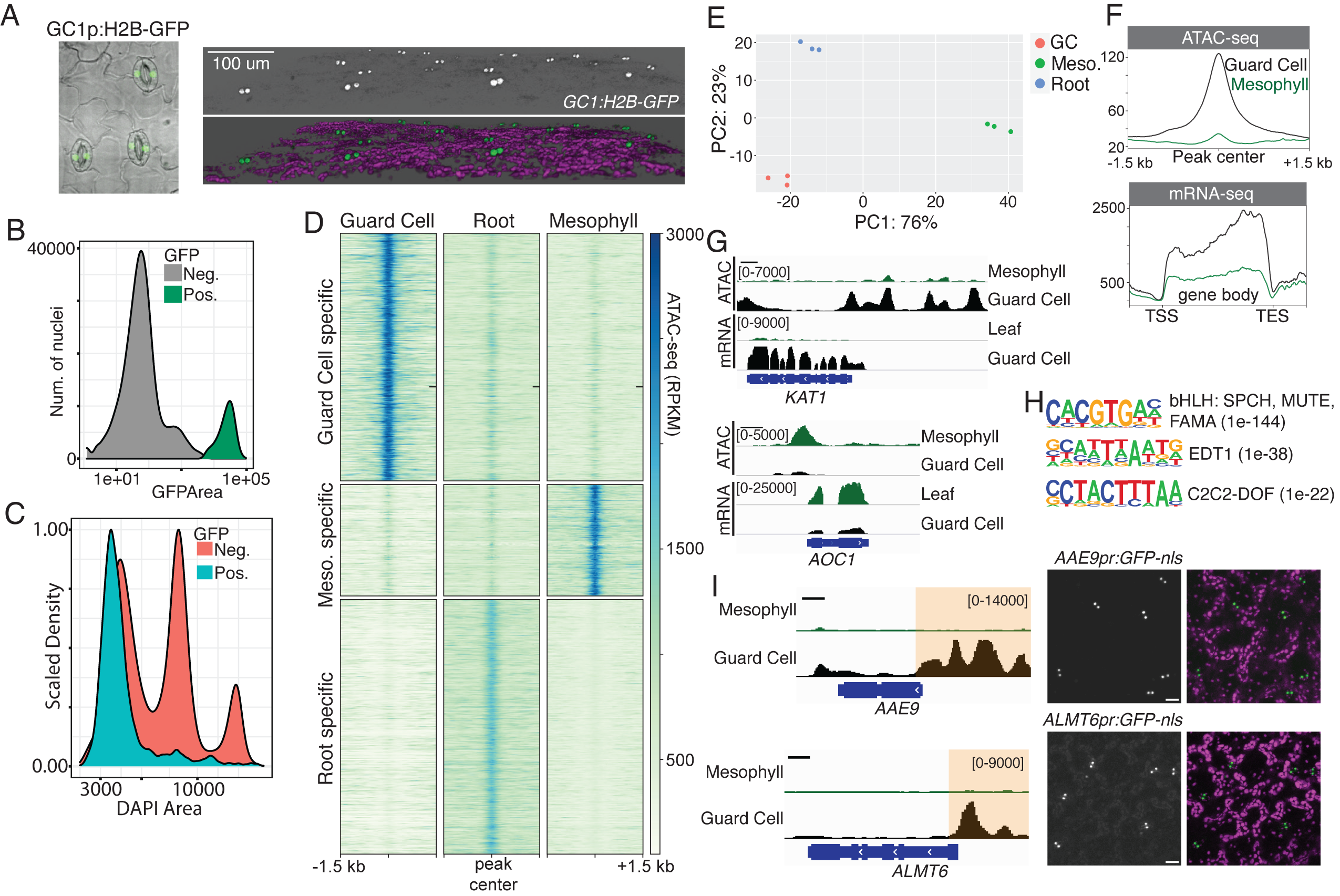
Cell-type specific epigenomics identifies guard cell specific regulatory elements. **A)** Left panel is a fluorescence micrograph showing guard cell specific expression of H2B-GFP. Right panel shows a 3D reconstruction from confocal imaging of a 5-week-old *GC1p:H2B-GFP* leaf (merged image shows GFP fluorescence in green and chlorophyll fluorescence in magenta). **B)** FACS analysis of nuclei (DAPI positive) from *GC1p:H2b-GFP* plants. A GFP density plot defines two distinct populations separated by GFP intensity. **C)** DAPI density plot shows that GFP positive nuclei (blue population) are primarily diploid, while GFP negative nuclei (pink population) are distributed over multiple ploidies. **D)** Chromatin accessibility heatmaps comparing ATAC-seq signal (RPKM-normalized) in guard cell, mesophyll, or root cell nuclei. ATAC-seq peaks are separated into guard cell enriched (3272 ACRs), mesophyll enriched (1472 ACRs), and root enriched (3364 ACRs) sets defined by differential accessibility (p.adj < 0.001, Fold-Change > 2). **E)** Principal component analysis (PCA) comparing ATAC-seq biological replicates derived from guard cell, mesophyll, or root nuclei. **F)** A peak-centered ATAC-seq metaplot and a gene-body centered RNA-seq metaplot showing average chromatin accessibility at guard cell specific peaks and average transcript levels of nearest downstream genes (ATAC-seq peak within -2.5 kb and +0.5 kb of transcription start site). Guard cell derived signal is shown in black while mesophyll/leaf signal is shown in green. **G)** Genome browser images displaying ATAC-seq and RNA-seq signal for the indicated tracks near the guard cell expressed gene *KAT1* and the mesophyll cell expressed gene *AOC1*. Scale bars indicate 500 bp. **H)** Results of de novo motif analysis using homer are shown along with the best matching known TFs and the enrichment p-values. **I)** Transcriptional reporter lines generated using cloned guard cell specific ACRs upstream of the genes *AAE9* and *ALMT6,* the positions of the cloned sequences are indicated with orange highlighting on the genome browser images (scale bars indicate 500 bp). Confocal micrographs showing GFP fluorescence (left panel) or merged GFP (green) and chlorophyll (magenta) fluorescence (right panel). Scale bar represents 20 µm.

In parallel, we generated ATAC-seq samples from root nuclei and from FANS-isolated mesophyll nuclei labeled by expression of H2B-GFP from the promoter of the mesophyll expressed Rubisco small subunit 2B gene (Fig. S3A-C) (6). For each tissue/cell-type we compared ATAC-seq libraries derived from three biological replicates. We identified 30,985 ATAC-seq peaks across all samples (Dataset S2) and called cell-type enriched peaks using pairwise differentially accessibility analysis. Although most ATAC-seq peaks did not differ significantly among cell types, we found 3272 guard cell enriched ACRs, 1472 mesophyll enriched ACRs, and 3364 root enriched ACRs (Dataset S2). Heatmaps centered over these regions highlight characteristics of cell/tissue specificity in the observed patterns of chromatin accessibility (Fig. 2D, Fig. S4A). Importantly, biological replicates derived from the same tissue/cell-type clustered together in principal component analysis (PCA) underscoring the robustness of this approach (Fig. 2E).

To evaluate the relationship between guard cell enriched chromatin accessibility and gene expression we performed RNA-seq on both whole leaves and guard cell-enriched samples. We found that regions with high chromatin accessibility specifically in guard cells were associated with elevated transcript levels from the adjacent downstream genes, indicating that we are identifying functional cis-regulatory elements (CREs) (Fig. 2F). Although the correlation between overall promoter chromatin accessibility and downstream transcript level was weak in guard cells (Pearson’s r = 0.19, Fig. S4B), we found that differentially accessible ACRs showed higher correlation with downstream gene differential expression (Pearson’s r = 0.55, Fig. S4C). Examination of individual gene loci like the guard cell expressed gene *KAT1* (46–48) illustrated the connection between increased upstream accessibility and increased transcript levels (Fig. 2G and Fig S4D). By contrast, the mesophyll expressed gene *AOC1* had high chromatin accessibility in mesophyll cells, but minimal upstream ATAC-seq signal in guard cells (Fig. 2G). TF binding motif analysis on guard cell enriched ACRs returned de novo motifs with strong similarity to motifs recognized by known stomatal lineage transcriptional regulators such as the bHLH proteins SPCH, MUTE, and FAMA as well as EDT1 and DOF-type TFs (49–53) (Fig. 2H and Dataset S7). In contrast, we find that WRKY-type TF binding motifs are enriched in regions that are significantly less accessible in guard cells compared to roots and mesophyll cells (Table S7). However, these motifs are recognized by large transcription factor families containing many transcription factors that are not specific to guard cells, for instance SPCH and MUTE are active only early during stomatal lineage development (49, 50). By connecting guard cell specific ACRs with downstream guard cell enriched transcripts we were able to define a set of 827 putative regulatory elements preferentially active in guard cells (Table S4). To further confirm that our approach identified functional cis-regulatory regions, we generated transcriptional reporter constructs expressing nuclear localized GFP under the control of the guard cell specific ACRs upstream of the malate-transporter encoding gene *ALMT6* (54) and the wax biosynthesis enzyme-encoding gene *AAE9* (55). Both reporter constructs produced guard cell specific GFP signal in transgenic *Arabidopsis* leaves (Fig. 2I). In summary, we have developed a robust protocol for isolating pure populations of guard cell nuclei allowing us to discover cis-regulatory regions that are active in mature guard cells.

### ABA induces persistent chromatin remodeling in guard cells during stomatal closure

We next asked whether epigenomic reprogramming occurs during ABA-induced stomatal closure. Soil grown 6-week-old plants were sprayed with 50 uM ABA or control solution. After 3 hours, we confirmed a stomatal response by infrared photography (Fig. 3A). To determine if our ABA treatment led to significant transcriptional changes we performed RNA-sequencing on both whole leaf samples (Fig. S5A) and guard cell enriched samples (Fig. 3B). We observed that ABA triggered the differential expression of thousands of transcripts in both whole leaves and guard cells (2142 upregulated and 1574 downregulated in guard cells). As expected, ABA-induced genes are enriched for the pathways “response to water deprivation” (p_value = 1.7e-22), “response to abscisic acid” (pvalue = 4.47e-18), and “response to salt stress (Fig. S5D). Although most of the transcriptional changes induced by ABA were shared between guard cells and whole leaves, we found that 22.7% (488) of ABA-induced transcripts and 42.7% (673) of ABA-repressed transcripts were specific to guard cells (Fig. S5C, Dataset S5). For instance, ABA-regulated expression of the major facilitator protein-encoding gene *AT2G37900* and the gene *AAE9* was only evident in guard cells (Fig. S5E).

**Figure 3.**
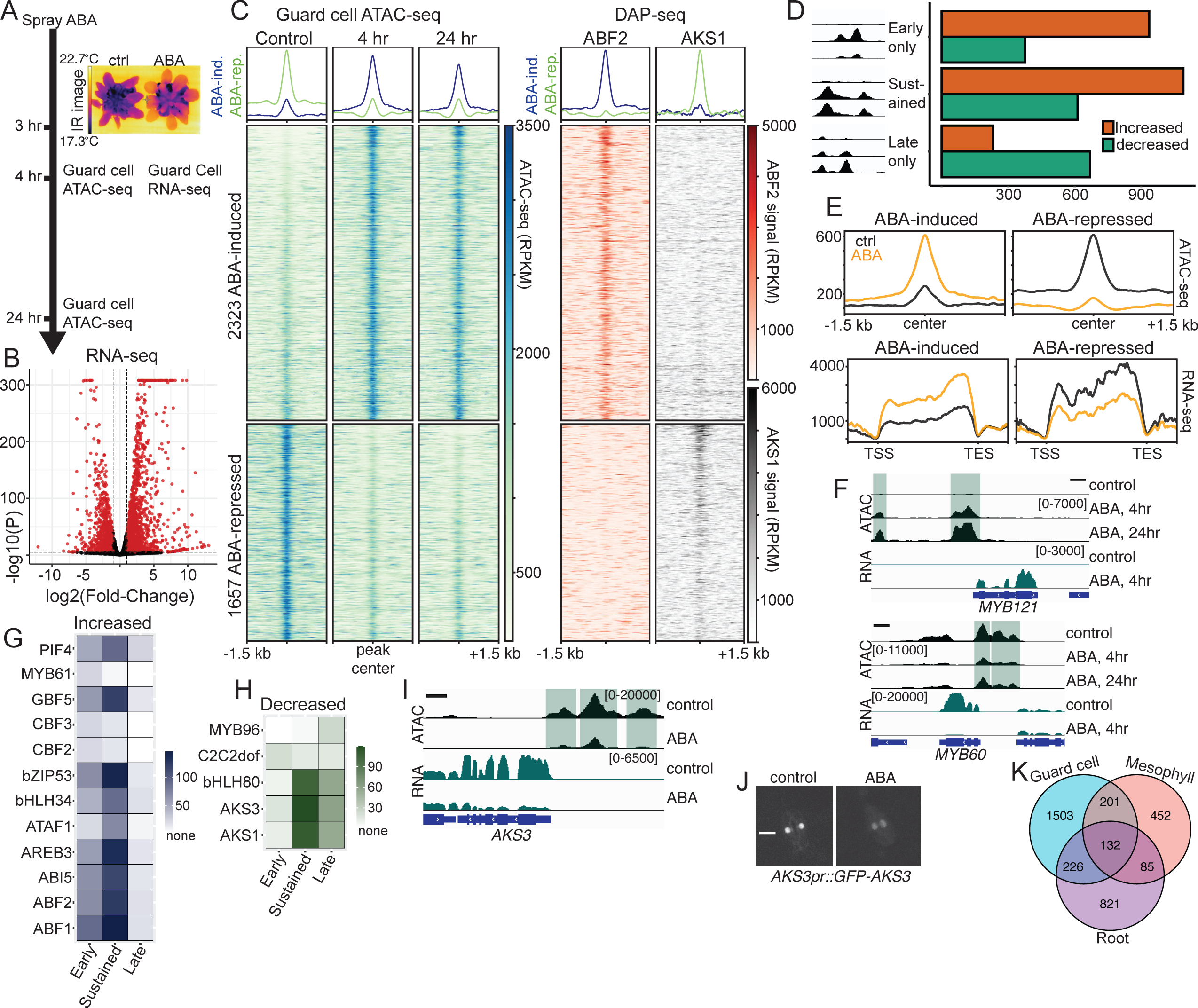
ABA triggers extensive and persistent chromatin remodeling in guard cells. **A)** Schematic showing the experimental outline. Plants were sprayed with ABA and infrared images were taken using a thermal camera to assess stomatal closure. Guard cell nuclei were purified at 4hrs and 24 hrs after treatment and used for ATAC-seq. **B)** Volcano plot of guard cell enriched RNA-seq data showing that ABA upregulates 2142 transcripts and downregulates 1574 transcripts (FDR < 0.001, FC > 2.0). Differentially expressed transcripts are colored in red. **C)** Heatmaps and profile plots of ATAC-seq signal (RPKM-normalized) in guard cell nuclei centered (+/- 1.5 kb) over 2323 regions with ABA-induced chromatin accessibility and 1657 regions with ABA-repressed chromatin accessibility (FDR < 0.001, FC > 2.0). The adjacent heatmaps show DAP-seq signal for ABF2 (in red) and AKS1 (in black) centered over the same ABA regulated ACRs, data from (39) and (56). **D)** Comparison of ABA-regulated peaks from 4hr and 24hr timepoints separated ACRs into three categories – early only, sustained (both early and late), and late only. The bar graph shows the number of ABA-increased and -decreased regions in each category. **E)** (Top) Metaplots showing average ATAC-seq signal centered over either ABA-induced or ABA-repressed ACRs and (bottom) RNA-seq metaplots showing average transcript level centered over gene bodies downstream of the same ACRs. Signal from ABA-treated plants is shown in orange and signal from control treated plants is shown in black. **F)** Genome browser images showing guard cell ATAC-seq and RNA-seq signal at a representative ABA-induced gene (*MYB121)* and an ABA-repressed gene (*MYB60*). Scale bars indicated 500 bp. **G,H)** Heatmaps showing TF binding motif enrichments (-log(p-value)) in ABA-increased **(G)** and ABA-decreased **(H)** ATAC-seq peaks. **I)** Genome browser images showing guard cell ATAC-seq and RNA-seq signal at the *AKS3* gene. Scale bars indicated 500 bp. **J)** Fluorescence micrograph showing expression of *AKS3pr::GFP-AKS3* in guard cells after treatment with control or ABA (4 hr). **K)** Overlap of ABA-induced accessible regions in guard cell nuclei, mesophyll nuclei, and root nuclei.

To evaluate if ABA changed chromatin accessibility and whether these changes are sustained over time following a single treatment, we isolated guard cell nuclei at 4 hours and 24 hours after spraying (Fig. 3A). We generated ATAC-seq libraries from three biological replicates per time point. PCA analysis showed that samples clustered together according to treatment (Fig. S6A). Differential chromatin accessibility analysis found that ABA extensively reshapes chromatin structure in guard cells with 2323 ACRs gaining accessibility and 1657 ACRs losing accessibility across the treatment time course (Fig. 3C, Fig. S7A and B, and Dataset S3). We next analyzed the temporal characteristics of this response by comparing the ATAC-seq datasets from the 4-hour and 24-hour timepoints. Our results allowed us to define three categories of ABA-regulated ACRs – early, sustained, and late (Fig. 3D). Strikingly, we found that 1093 of the 2079 (52.6%) ACRs opened by ABA at 4 hours remained open after 24 hours. Of the 975 ACRs closed by ABA at 4 hours 594 (61%) remained closed at 24 hours. Additionally, we found 230 ACRs gained and 682 ACRs lost accessibility only 24 hours after ABA treatment (Fig. 3D). Interestingly, we found differences in downstream gene GO term enrichment between early and late ACRs. For example, the terms “response to light” and “defense response” were strongly enriched in ABA-repressed ACRs only at 24 hours after ABA treatment (Table S6).

Consistent with our observations in seedling roots (Fig. 1D), we found that ABA-regulated ATAC-seq peaks in guard cells reside further away from transcription start sites (TSS) than non-regulated peaks (> 1kb), indicating that distal regulation may be a characteristic of ABA-controlled CREs (Fig. S6B). ABA-induced ACRs in guard cells are clearly associated with downstream genes that are upregulated transcriptionally by ABA, while ABA-repressed ACRs are associated with genes downregulated by ABA (Fig. 3E). Indeed, ABA-induced changes in chromatin accessibility were correlated with ABA-induced differential expression of downstream genes (r = 52, Fig. S6C). Examination of individual genes like the ABA-upregulated *MYB121* and the ABA-downregulated *MYB60* (56) illustrate the connection between upstream chromatin accessibility and transcript levels (Fig. 3F). Interestingly, we found that ABA modifies chromatin accessibility upstream of many known regulators of stomatal conductance including *KAT1*, *NCED3*, and *HT1* (57) as well as many other genes with no known stomatal functions (Fig. S6D-K, Table S3).

We next performed TF binding motif analysis on early, sustained, and late ABA-regulated ACRs using a set of DAP-seq derived motifs (43) restricted to those TFs with guard cell expression in our RNA-seq datasets. Both early and sustained ABA-induced ACRs had strong enrichment for motifs recognized by ABRE-binding TFs including ABI5, ABF1,2, and 3 (Fig. 3G). Interestingly, we found that the motifs recognized by CBF2, CBF3, and MYB61 (58–60) were preferentially enriched in early ABA-induced ACRs. ABA-repressed ACRs showed strong enrichment for motifs recognized by a clade of related bHLH-type TFs (Fig. S6L) including AKS1, AKS3, and bHLH80 in both sustained and late categories (Fig. 3H). In contrast, MYB96 motifs were enriched only in late appearing ABA-repressed ACRs (Fig. 3H). Collectively, our results demonstrate that ABA can induce persistent changes to chromatin in guard cells and expose the temporal regulation of different TFs during this response.

By analyzing published genome-wide DAP-seq datasets (39, *56*) we found strong ABF2 DAP-seq signal coincident with ABA-induced ACRs in guard cells (Fig. 3C). DAP-seq signals for ABF1, ABF3, and ABF4 were also enriched over regions that gain accessibility in response to ABA, although to a lesser extent than ABF2 (Fig. S6M). In contrast, ABA-repressed ACRs contained AKS1 DAP-seq binding sites (Fig. 3C). Additionally, unbiased de novo motif discovery (62) returned motifs similar to ABF1 and AKS3 binding sites as the most enriched in ABA-activated and ABA-repressed ACRs, respectively (p = 1e-119 and p = 1e-57). Metaplots of ATAC-seq signal centered over peaks containing these motifs further supported our observation that ABA elevates chromatin accessibility at ABF binding sites and represses chromatin accessibility at AKS binding sites (Fig. S6N). Interestingly, ABA closed chromatin upstream of *AKS3* and decreased *AKS3* transcript levels in guard cells (Fig. 3I). We also observed that GFP-AKS3 signal decreased in ABA-treated guard cells when expressed from its native promoter (Fig. 3J).

Furthermore, by comparing ABA-induced ACRs from guard cells, mesophyll cells (Fig. S3E-G), and root cells (Fig. 1) we uncovered significant tissue/cell-type specificity in this response (Fig. 3K, Fig S7C-L). Interesting, we found that ABA had a much larger impact on chromatin accessibility in guard cells than in the other cell types with more ABA regulated regions (2079 ACRs compared with 1293 in roots and 926 in mesophyll) and larger fold changes in accessibility (Dataset S3). In conclusion, we show that ABA drives large scale changes to chromatin structure in guard cells and implicate major roles for the activation of ABFs and inhibition of AKSs in this response.

### ABF transcription factors are required for ABA-induced chromatin opening in guard cells

The mechanisms that cause ABA-specific chromatin opening are unknown. Given the strong enrichment of ABF binding motifs in ABA-activated ACRs, and their coincidence with ABF *in vitro* binding sites, we wondered if these TFs might be required for reshaping chromatin during ABA-induced stomatal closure. To test this, we used the *abf1/2/3/4* (*abfx4*) quadruple mutant which lacks four related ABF proteins that are prominent during vegetative growth (25, 26). We first performed RNA-seq following ABA treatment on leaves from 6-week-old *abf1/2/3/4* (*abfx4*) mutant plants (Fig. S8A). Of the 2257 transcripts that are upregulated by ABA in wild type leaves, 1897 (∼84%) depend on ABF1/2/3/4 (Fig. S8B). Interestingly, although ABA increased *ABF1/2/3/4* transcript levels in guard cells we did not find that ABA altered chromatin accessibility in the upstream regions near the corresponding genes (Fig. S8C). We conclude that ABF1/2/3/4 are required for the bulk of ABA-induced gene expression in mature leaf tissue. Plants lacking *ABF1/2/3/4* (*abfx4*) are highly sensitive to drought stress and lose water faster than wild type plants in detached leaf assays (26). Consistent with this, we observed that *abfx4* mutant plants exhibited higher steady-state stomatal conductances than *Col-0* control plants (Fig. S8D).

Under non-stressed conditions, guard cells have a higher ABA concentration than other leaf cell types (42, 63). Recent studies have provided evidence for basal ABA signaling in guard cells and defined important roles for basal ABA in stomatal function (64, 65). By generating guard cell FANS lines in an *abfx4* mutant background (Fig. S9A) we were able to assess the impact of the loss of ABF1/2/3/4 on guard cell chromatin in non-stressed plants. In the absence of ABA treatment, we observed minimal differences in genome-wide patterns of chromatin accessibility between *WT* and *abfx4* guard cells (Fig S9B, C and Dataset S3) suggesting that basal ABA signaling in guard cells may regulate mainly post-transcriptional stomatal closing mechanisms but does not significantly regulate chromatin structure.

Next, we asked if the ABF1/2/3/4 are required for the ability of ABA to reshape chromatin structure. Strikingly, loss of *abf1/2/3/4* strongly impaired ABA-induced chromatin opening in guard cells but had less impact on ABA-triggered chromatin closing (Fig. 4A, Dataset S7). Of the 2079 ABA-induced ACRs, 1539 (∼74%) required ABF1/2/3/4 for gained accessibility (Fig. 4B). ABF-dependent ABA-induced ACRs were strongly enriched for ABREs (example ABF1, p = 1e-86), while ABF-independent ABA-induced ACRs were most strongly enriched for CBF1/2/3 motifs (example CBF1, p = 1e-16) (Dataset S7). Examination of individual loci like the ABA-activated genes *DTX37* and *HSFA6B* revealed a close connection between abf-dependent increases in upstream chromatin accessibility and abf-dependent transcription (Fig. 4C).

**Fig 4.**
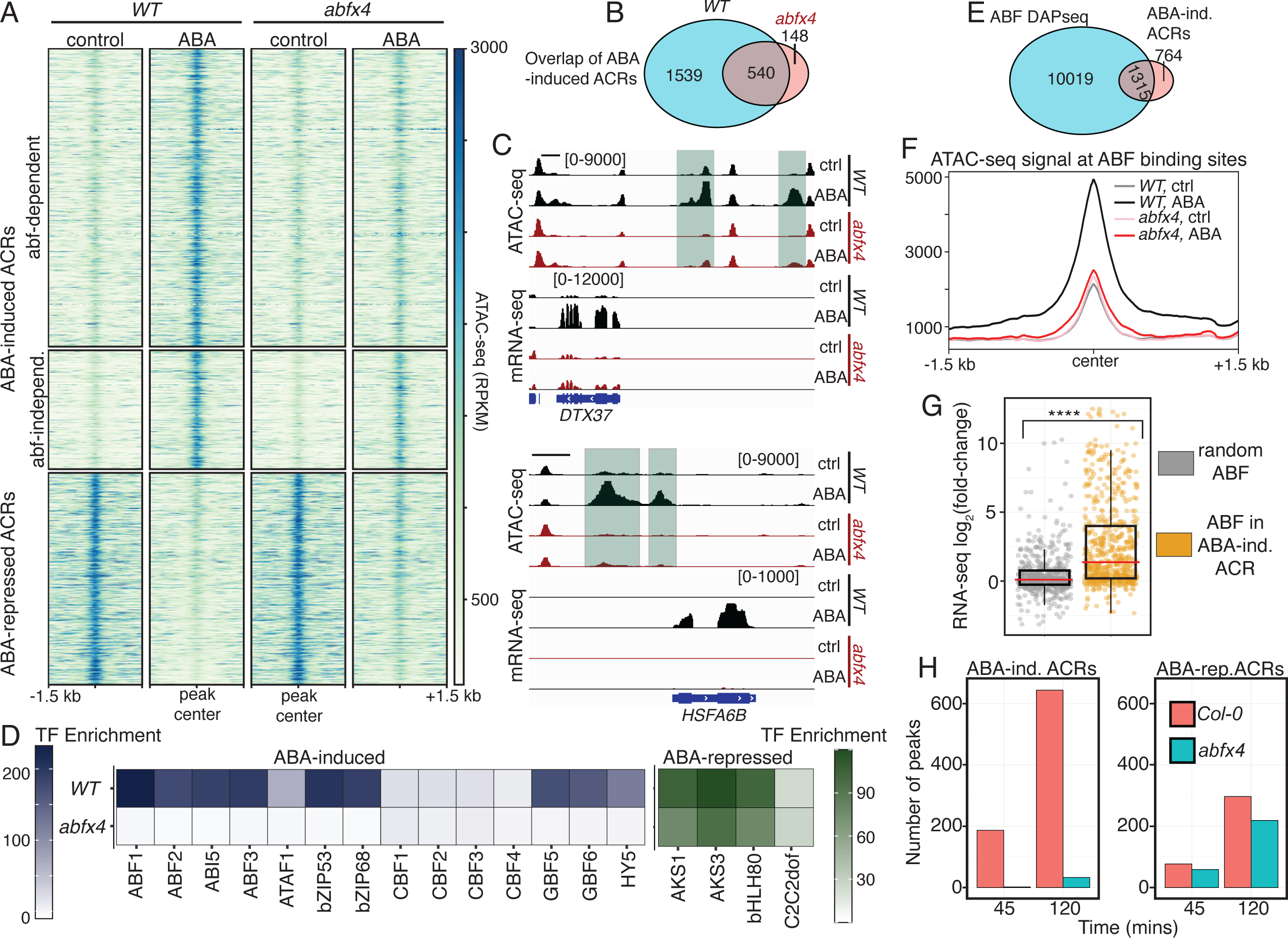
ABF transcription factors are required for ABA-induced chromatin opening in guard cells. **A)** Heatmaps of ATAC-seq signal (RPKM-normalized) in either *WT* or *abfx4* mutant guard cells centered over 2079 regions with ABA-induced accessibility and 975 regions with ABA-repressed accessibility (FDR < 0.001 and FC > 2.0). ACRs are subdivided by requirement for ABF1/2/3/4 **B)** Overlap of ABA-induced ATAC-seq peaks in *WT* guard cells with those in *abfx4* mutant guard cells. **C)** Genome browser snapshots showing ATAC-seq and RNA-seq signal near two ABA-induced genes *DTX37* and *HSFA6B*. Scale bars indicate 1 kb. **D)** Enrichment (-log(p-value)) of the indicated transcription factor binding motifs in ABA regulated ATAC-seq peaks in either *WT* or *abfx4* mutant guard cell nuclei. **E)** Overlap between ABF DAP-seq peaks and ABA-induced accessible chromatin regions (ACRs). **F)** Peak-centered metaplot showing ATAC-seq signal in the indicated samples at ABA-induced ACRs containing ABF DAP-seq binding sites. **G)** 1000 ABF DAP-seq peaks overlapping ABA-induced ATAC-seq peaks or 1000 random ABF DAP-seq peaks were assigned to nearest downstream genes (within -2.5 kb and +0.5 kb of TSS). The plot shows the ABA-induced fold changes within these two sets of downstream transcripts. Asterisks (****) indicate that the distributions are significantly different by Welch’s t-test (p < 1E-7) and the red bars represent the means of the distributions. **H)** Bar plots representing the number of ABA-activated or ABA-repressed ACRs over time in *Col-0* or *abfx4* mutant roots.

We next performed TF motif analysis on ABA-regulated accessible chromatin regions in both *WT* and *abfx4* mutant guard cells. As expected, *abfx4* mutants were depleted for ABRE motif enrichment in ABA-induced ACRs but notably retained enrichment for AKS1, AKS3, and bHLH80 motifs in ABA-repressed ACRs (Fig. 4D, Dataset S7). Additionally, although most TF binding sites lost enrichment in ABA-induced ACRs in the absence of ABF1/2/3/4, we found that enrichment for motifs bound by CBF1/2/3 was preserved. TF binding motifs are often information poor and the presence of a motif at a genomic location does not always lead to TF action. Analysis of published ABF1/2/3/4 binding assays conducted on naked genomic DNA (61) uncovered 11334 binding sites, but only 1315 resided within ABA-induced ACRs in guard cells (Fig. 4E). However, ABA-induced chromatin opening at these ACRs containing ABF-binding sites was strongly impaired in *abfx4* mutant guard cells (Fig. 4F). We reasoned that chromatin structure may determine which *in vitro* ABF binding sites are functional. To test this idea, we examined if ABF DAP-seq peaks found within ABA-induced ACRs could predict ABA-activated transcription. Using our guard cell RNA-seq data, we found that ABF binding sites that reside within regions of ABA regulated open chromatin were more strongly associated with downstream ABA-induced genes than were an equal number of random ABF-binding sites (Fig. 4G).

To examine the temporal characteristics of ABF-dependent chromatin remodeling we focused on roots where ABA can reshape chromatin by 45 mins (Fig. 1C). We performed ATAC-seq on isolated *abfx4* mutant root nuclei over an ABA treatment time course and found that ABA fails to drive progressive chromatin opening in this background (Fig. S10). In *abfx4* mutant roots, 45 mins of ABA treatment induced only 2 ACRs compared to 163 in wild type roots. By 2 hours of ABA treatment, only 27 ABA-induced ACRs were present in *abfx4* mutants compared with 601 in wild type (Fig. 4H, Fig. S10A, and Dataset S3). However, ABA repressed a similar number of root ACRs in the absence of ABF1/2/3/4 (Fig. 4H and Fig. S10A). Although our data support a general requirement for ABFs in ABA-induced chromatin opening in roots and guard cells, we do find that some guard cell specific ABA-induced ACRs are ABF-dependent (Fig. S10D,E). Collectively, these results demonstrate that the ABF1/2/3/4 transcription factors are required for triggering genome-wide chromatin opening in response to ABA.

### ABA and changes to atmospheric CO_2_ concentration induce distinct chromatin remodeling programs in guard cells

Because we observed that ABA has extensive effects on chromatin accessibility in guard cells, we wondered whether chromatin structure may respond to other stimuli that drive stomatal movements. Here we focus on changes to atmospheric CO_2_ concentration. Exposure of plants to low CO_2_ stimulates stomatal opening while high CO_2_ drives stomatal closure (Fig 5A). The exact role of ABA signaling during CO_2_ induced stomatal responses is unclear and is a matter of some debate (64–66). Recent studies suggest that although high-CO_2_ induced stomatal closure requires basal ABA signaling it does not involve further SnRK2 kinase activation (64, 65). Comparing how these signals impact genome activity could provide new insight into whether and how the mechanisms underlying guard cell responses to ABA and CO_2_ may differ.

**Fig 5.**
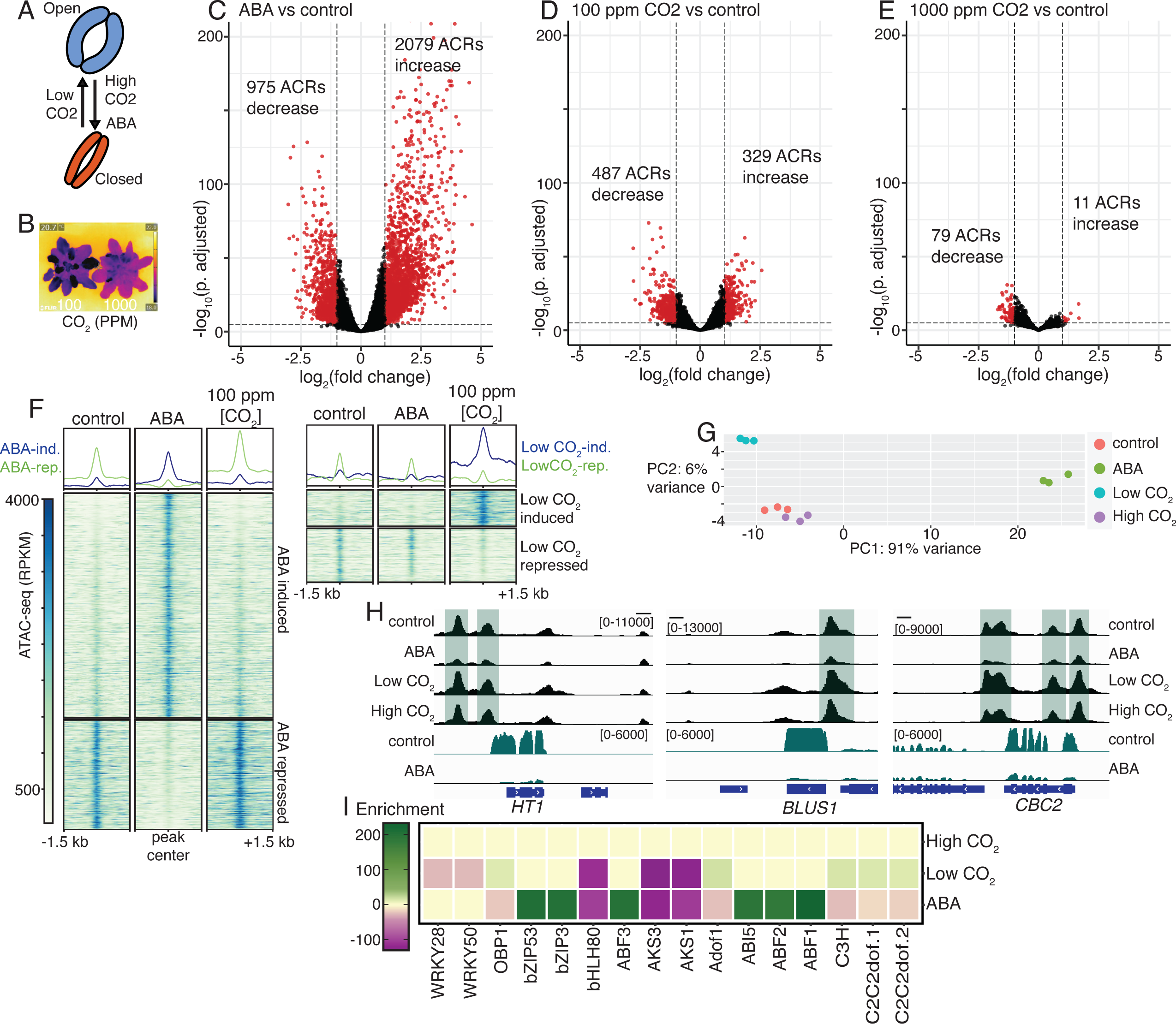
ABA and changes in CO_2_ concentration induce distinct chromatin remodeling programs in guard cells. **A)** Diagram showing the effects of different stimuli on the stomatal aperture. **B)** Representative IR image showing differences in leaf temperature due to CO_2_ induced stomatal movement. **C-E)** Volcano plots of differential ATAC-seq results showing **C)** ABA-regulated chromatin accessibility, **D)** low CO_2_ regulated chromatin accessibility, and **E)** high CO2 regulated chromatin accessibility. The numbers of differentially accessible peaks (colored in red, FDR < 0.001 and FC > 2.0) in each condition are indicated on each plot. **F)** Heatmaps and profile plots of guard cell ATAC-seq signal centered over either ABA-regulated ACRs (2079 induced and 975 repressed) or low CO_2_-regulated ACRs (329 induced and 487 repressed). Libraries are derived from plants exposed to the indicated treatments. **G)** Principal component analysis (PCA) shows that ABA, low CO_2_, and high CO_2_ introduce separate variation to genome-wide chromatin accessibility. **H)** Genome browser images showing ABA-repression of chromatin accessibility upstream of *HT1*, *BLUS1*, and *CBC1* – three positive regulators of low CO_2_ induced stomatal opening. Scale bars indicate 1 kb. **I)** Enrichment (-log P_value) of the indicated transcription factor binding motifs in ABA, Low CO_2_, or High CO_2_ regulated regions. Negative enrichment scores indicate motifs overrepresented in treatment-repressed regions. No significant enrichment for any of these motifs was found in High CO_2_ regulated ACRs.

To generate low and high CO_2_ guard cell ATAC-seq libraries we exposed 6-week-old plants to either 100 ppm CO_2_ or 1000 ppm CO_2_ for 4 hrs, two treatments that induce robust stomatal responses (Fig 5B). Surprisingly, in contrast to the extensive effect of ABA on chromatin accessibility in guard cells (Fig. 5C), we observed smaller changes in response to low CO_2_ (Fig. 5D), and minimal changes following treatment with high CO_2_ (Fig. 5E). Following exposure of plants to 100 ppm CO_2_, 329 ACRs gained accessibility and 487 ACRs lost accessibility while treatment with 1000 ppm CO_2_ induced only 11 ACRs and repressed 79 ACRs (Dataset S3). Heatmaps of ATAC-seq signal at ABA and Low CO_2_-regulated regions highlighted the specificity of these responses (Fig. 5F, Fig. S7A,B). PCA results comparing our guard cell ATAC-seq datasets from different conditions further support these findings and revealed little difference between control and high CO_2_ treatments (Fig. 5G). Examination of individual loci highlights the specificity of these responses, for instance chromatin opening upstream of *ERF113* only occurred in response to ABA while the upstream region of the PP2C-encoding *AT2G25070* gains accessibility only in response to low CO_2_ (Fig. S11B-E).

Analysis of the set relationships among ABA, low CO_2_, and high CO_2_ regulated ACRs revealed little overlap suggesting that these stimuli control distinct cis-regulatory elements in the genome (Fig. S11A). We did find overlap between ABA-induced and low CO_2_-repressed ACRs and between ABA-repressed and low CO_2_-induced ACRs. ABA-repressed regions had elevated average ATAC-seq signal in guard cells isolated from plants exposed to 100 ppm [CO_2_] (Fig. 5F). For instance, ABA-repressed chromatin accessibility upstream of *ALMT6* while low-CO_2_ treatment enhanced accessibility (Fig. S11D). Interestingly, ABA repressed chromatin accessibility upstream of *HT1*, *BLUS1*, and *CBC2* (67), all key positive regulators of low CO_2_- and blue light-induced stomatal opening, while high CO_2_ had no effect (Fig. 5H). These data indicate that chromatin remodeling represents an important cross talk point for these stimuli, ensuring robust stomatal closing or opening, depending on the physiological conditions. DNA binding motif analysis further emphasized the distinct regulatory impacts of changes in CO_2_ concentration and ABA on the genome (Table S7). In contrast with ABA-induced ACRs, we did not find enrichment of ABF binding motifs in low or high CO_2_ regulated chromatin (Fig. 5I). Instead, low CO_2_ gained ACRs were enriched for motifs recognized by C2C2dof-type TFs which were oppositely enriched only in ABA-repressed ACRs. Collectively, our data demonstrate how different signals that control stomatal movement deploy distinct gene regulatory programs in guard cells.

## DISCUSSION

Abscisic acid triggers changes in gene expression, but whether this transcriptional response depends on chromatin remodeling remained unknown (5). Here, we reveal massive ABA-induced chromatin remodeling pointing to an orchestrated reprogramming of the *Arabidopsis* epigenome by specific sets of transcription factors. We identify regulatory regions of the genome specifically active in guard cells and uncover how the accessibility of these sequences is modified by signals that control stomatal aperture size.

Guard cell regulation of stomatal aperture size is essential for plant survival under drought stress. Such regulation requires that the guard cells forming the stomatal pore integrate numerous physiological and environmental cues. In this study, we profiled genome-wide chromatin accessibility in tens of thousands of guard cells isolated from soil-grown plants exposed to different stimuli that trigger stomatal movements. Our results lead us to propose that guard cell ABA signaling controls the balance between two opposing chromatin states. Under unstressed conditions, AKS proteins maintain open chromatin at upstream regulatory regions of ABA-repressed genes while ABA-activated genes are kept silent by closed chromatin. Under abiotic stress such as drought, ABA levels rise in guard cells thereby activating SnRK2 kinases which subsequently inhibit AKSs and activate ABFs. This leads to chromatin opening upstream of abiotic stress-activated genes and the loss of chromatin accessibility upstream of stress-repressed genes.

We uncovered patterns of chromatin accessibility specific to guard cells (Fig. 2D) and identified cis-regulatory regions supporting guard cell specific gene expression (Fig. 2F-I and Dataset S4). We found that ABA induces extensive and cell-type specific chromatin remodeling and that guard cells possess a more extensive response than mesophyll and root cells (Fig. 3K and Fig. S7C-L). Importantly, we find that these ABA-induced chromatin remodeling events correlate with ABA-regulated gene expression (Fig. 3E and S6C). By performing TF-binding motif enrichment and analyzing published DAP-seq data, we hypothesized that the bZIP transcription factors ABF1/2/3/4 may function in ABA-induced chromatin remodeling in guard cells (Fig. 3C,G and Fig. S6M). Interestingly, we find that among *in vitro* ABF binding sites, those occurring within ABA-opened chromatin are most strongly associated with ABA-induced transcription (Fig. 4G), indicating that regulated chromatin accessibility may determine which genomic ABF binding sites are functional. Our experiments in *abf1/2/3/4* quadruple mutant guard cells demonstrate that these four transcription factors are essential for the majority of ABA-triggered chromatin opening (Fig. 4). Furthermore, experiments in root nuclei show that in *abf1/2/3/4* quadruple mutants, even early ABA-induced chromatin remodeling events are strongly impaired (Fig. 4H and Fig. S10), suggesting that ABFs initiate chromatin opening. How ABFs accomplish chromatin opening remains to be determined, but existing literature could be interpreted to support two possible and not necessarily mutually exclusive explanations. (1) Although nucleosomes can restrict TF binding to DNA, some TFs, known as pioneer factors (68, 69), possess the ability to bind their target sequences even within inaccessible chromatin regions. At present, only a handful of pioneer factors have been documented in plants (70). While our data are consistent with ABFs acting as pioneer factors, future biochemical experiments reconstituting ABF binding to nucleosomal DNA are needed to investigate this model. (2) ABFs may recruit chromatin modifying proteins to target sequences which could increase chromatin accessibility. Interestingly, several such molecules including the SWI/SNF ATPase BRM (71, 72) and the histone deacetylase HDA9 (73) interact with known ABA signaling components. Furthermore, ABF-binding proteins known as AFPs, which negatively regulate ABA signaling, have been shown to physically interact with histone and chromatin modifying factors (74).

Although our data support a general requirement for ABF proteins in both roots and guard cells, we also observe that ABA alters chromatin accessibility in a cell-type specific manner. At present we do not understand how ABFs might contribute to cell-type specific regulation of chromatin. Beyond ABF1/2/3/4, our analyses implicated additional TFs in ABA-dependent genome reprogramming (Fig. 3F and 3G), which is consistent with prior research showing that ABA regulates the binding of multiple transcription factors (75). We observed that ∼26% of ABA-gained ACRs (540 ACRs) still open in *abfx4* mutant guard cells in response to ABA (Fig. 4B). These ABF-independent regions retained enrichment for CBF1/2/3 binding motifs, suggesting that CBF transcription factors display ABF-independent binding (Fig. 4D). Finally, although ABA-induced chromatin opening is severely impaired in *abf1/2/3/4* quadruple mutant guard cells, we cannot exclude the possibility that other ABRE-binding TFs, in particular ABI5 (76, 77), participate in this process as well.

Notably, our data suggest that the related bHLH transcription factors AKS1/2/3 play a larger role in the regulation of ABA-repressed genes than was previously thought. Prior research had shown that SnRK2 kinase-dependent inhibition of AKS1/2 was required for the ABA-dependent transcriptional repression of the K+ ion channel gene *KAT1* in guard cells (24). Surprisingly, we found that AKS1 bound many regions in the genome that lose chromatin accessibility in guard cells following ABA treatment (Fig. 3C). Additionally, motifs recognized by AKS1/3 and the related protein bHLH80 were enriched in ABA-repressed ACRs in guard cells (Fig. 3H), as well as in root and mesophyll cells (Fig. 1F, Fig. S3G, Table S7). This observation was further supported by de novo motif analysis (Fig S6N). AKS1 and AKS2 are inhibited by direct ABA-dependent SnRK2 phosphorylation, but their expression was not controlled by ABA signaling (44). We found that in guard cells, ABA represses *AKS3* expression implying that ABA-dependent inhibition of AKS proteins occurs through both transcriptional and phosphorylation-dependent mechanisms (Fig. 3I and J).

Elevation in the CO_2_ concentration in leaves triggers rapid stomatal closing (34). The extent to which elevated atmospheric CO_2_ signals through the ABA signal transduction pathway is currently under debate. Different research groups using the same mutants have come to different conclusions regarding direct CO_2_ signaling via the early ABA receptor/SnRK2 kinase pathway during CO_2_-induced stomatal closure. Our results exposed major differences in how guard cells interpret these stimuli. In contrast to the extensive and persistent changes to chromatin structure induced by ABA (Fig. 3), exposure to elevated CO_2_ had minimal impact on genome-wide chromatin accessibility in guard cells (Fig. 5). While ABA controlled chromatin accessibility at 3018 ACRs in the genome, we found that only 90 ACRs were regulated by elevated CO_2_ (Fig 5C-E). Additionally, the strong enrichments of ABF binding motifs in gained ACRs and AKS binding motifs in lost ACRs, evident following ABA treatment, are absent during the response to elevated CO_2_ (Fig. 5I). The disparity between these two signals at the level of chromatin mirrors the disparity between characteristics of their respective physiological responses. Unlike ABA- and drought-triggered stomatal closure, elevated CO_2_-induced stomatal closure is rapidly reversed upon return to ambient CO_2_ (64). Such rapid changes in the intercellular CO_2_ concentration in leaves occur for example with alternating light intensities (e.g. passing cloud cover) and require rapid reversals of stomatal CO_2_ responses. Furthermore, we found that exposure to low CO_2_ triggers a distinct program of chromatin remodeling in guard cells (Fig. 5F-I).

Finally, our results may open new avenues for investigation into two poorly understood features of stomatal biology. Studies in multiple plant species have documented a phenomenon referred to as stomatal “memory” (78, 79). Following drought stress and re-watering, plants typically exhibit delayed and/or slow re-opening of stomata despite rapid recovery of plant water status. Furthermore, guard cells display transcriptional memory where repetitive dehydration stress leads to altered transcript levels in subsequent stress events (80–82). We found that ABA triggered changes to chromatin accessibility in guard cells that can persist for up to 24 hours, which could prime the genome for subsequent abiotic stress exposure (Fig. 3). These shifts occurred at thousands of locations in the genome including upstream of many genes known to regulate stomatal movements as well as many genes with potential functions in stomatal physiology that could be the basis for future studies (Fig. S6 and Dataset S3). Moreover, our findings support crosstalk between ABA and low CO_2_ signaling at the chromatin remodeling level. For example, we uncovered that ABA, but not high CO_2_, triggered chromatin closing upstream of genes encoding key CO_2_-sensing and signaling proteins that promote low CO_2_-induced stomatal opening, including *HT1*, *BLUS1*, and *CBC2* (Fig. 5H). This response may contribute to the long-term stomatal closing triggered by drought and ABA by inhibiting mechanisms for low CO_2_-induced stomatal opening. ABA-triggered chromatin dynamics may provide a basis for long-term reprogramming of the guard cell genome for abiotic stress survival. We envision that these sustained changes to chromatin structure may store the experience of drought stress and contribute to long-term adjustments of stomatal function.

## METHODS

### Plant material and growth conditions

The Columbia-0 (Col-0) accession of *Arabidopsis thaliana* was used as the wild-type background. Experiments on seedlings were conducted using plants germinated and grown on 1/2 Murashige and Skoog (MS) media (pH 5.7-8) solidified with 1% Phyto-agar. Experiments on mature plants were conducted using plants grown in soil (Sunshine Mix #1) filled plastic pots incubated in a growth chamber (Conviron) under a 12hr-12hr light-dark cycle, a light intensity of 100 µmol m^-2^ s^-1^, a temperature of 22 C, ambient [CO_2_], and a relative humidity of 65%. To minimize the impact of environmental variation all experiments were performed on plants grown in two identical growth chambers equipped with [CO_2_] control.

ABA treatments in plate-based experiments were performed using a mesh (100 µm polyester) transfer method. Seeds were germinated on ½ MS agar plates overlaid with sterilized polyester mesh (100 µm). ABA treatments were initiated by transferring 8-day-old seedlings in mass to plates containing 50 µM ABA (Sigma cis/trans-abscisic acid) or to plates containing vehicle control (Ethanol). ABA treatments on soil grown plants were performed by spraying plants with 50 µM ABA diluted with 0.01% Silwet (in water) or with vehicle control. To assess treatment responses, leaf temperature was measured using an infrared thermal imaging camera (T650sc; FLIR).

### Plasmid DNA and plant transformation

Constructs for cell-type specific nuclear sorting were prepared using multi-site gateway technology (Invitrogen). The ORF encoding the histone H2B (*AT5G22880*) was cloned in a pDONR221 entry vector. The cell type specific promotors RBCp and GC1p, from the Rubisco small subunit 2B gene (*AT5G38420)* and the *GASA9* (*AT1G22690*) gene respectively, were cloned from the *Arabidopsis* genome into the gateway entry vector pDONR-p4p1r. Final plant transformation constructs were generated by performing multisite gateway recombination into the pP7m34GW destination vector. The plasmid for expressing *GFP-AKS3* under the *AKS3* (*AT2G42280*) endogenous promotor was generated using GreenGate cloning (83). The cloned *AKS3* upstream regulatory sequence was defined using guard cell ATAC-seq data. Transcriptional reporter constructs expressing nuclear localized GFP under the guard cell specific ACRs upstream of *AAE9* (*AT1G21540*) and *ALMT6* (*AT2G17470*) were generated using GreenGate cloning. Primer sequences are available in Dataset S8. Constructs were transformed into *A. thaliana* Col-0 plants using floral dipping with *Agrobacterium tumefaciens* strain GV3101.

### Fluorescence-activated nuclear sorting (FANS)

FANS on seedling/root nuclei and mesophyll nuclei was performed similar to (40) with small modifications. Briefly, whole seedlings, surgically isolated roots, or leaves were chopped vigorously with a clean razor blade in 2-3 ml of ice-cold FANS-lysis buffer (15 mM Tris-HCl pH7.5, 20 mM NaCl, 80 mM KCl, 0.2 mM spermine, 5 mM 2-ME, 0.5 mM spermidine, 0.2% IGEPAL CA-630, Roche mini EDTA-free Protease Inhibitor Cocktail) for 5 minutes in a cold glass petri dish. The subsequent lysate was filtered through a 40 µm filter unit before transferring to a 15 ml tube containing 3 ml of FANS-Sucrose buffer (20 mM Tris-HCl pH 8.0, 2 mM MgCl_2_, 2 mM EDTA, 15 mM 2-ME, 1.7 M sucrose, 0.2% IGEPAL CA-630). Nuclei were pelleted at 2500xG at 4°C for 25 mins, after which the supernatant was carefully removed. The nuclei pellet was then gently resuspended, using clipped pipette tips, with 600 µl FANS-lysis buffer supplemented with DAPI, and then immediately used for flow cytometry.

For experiments on guard cells, the leaves of ∼40 six-week-old plants were harvested and blended in 500 ml of ice-cold water for 2×40 secs with 1 min between pulses. The blended sample was poured over a 100 µm mesh to remove mesophyll cells and collect epidermal enriched fragments. The mesh was rinsed with ∼200 ml of cold water and gently blotted with paper towels to remove excess liquid. Thin sections of epidermal tissue were then flash-frozen in liquid nitrogen. Epidermal sections were ground to a fine powder with a frozen mortar and pestle. After grinding, powder was transferred to a new mortar with a frozen metal spatula. Immediately 12-15 ml of ice-cold FANS-lysis buffer was added, and the powder was gently resuspended with a clean pestle. The subsequent lysate was filtered through a 40 µm filter unit before transferring to a 15 ml tube containing 6 ml of FANS-Sucrose buffer. Nuclei were then pelleted and prepared for FACS as above.

Flow cytometry was conducted on a BD FACS Aria II using a 70 µm nozzle with a flow rate < 2 and the following photomultiplier parameters: Forward Scatter (FSC) – 250 eV, Side Scatter (SSC) – 220 eV, BV421 (DAPI channel) – 550 eV, FITC (GFP channel) – 340 eV. Sort gates were defined empirically using the FSC and SSC channels to exclude debris and potential doublets. Before sorting GFP labeled nuclei, FACS analysis on unlabeled control samples (Col-0 nuclei) was performed to define background fluorescence signal. When isolating guard cell nuclei, sort gates were established to purify exclusively 2N (diploid) and GFP-positive nuclei. To evaluate the purity of sorted samples, a fraction (∼10% of total volume) was re-run under the same sorting program (Fig. S2D). Sorted nuclei were collected in 600 µl of FANS-lysis buffer in a refrigerated tube chamber. After sorting, nuclei were pelleted by centrifugation at 1000xg for 10 mins at 4C. Nuclei pellets were gently washed with 1 ml of 10 mM Tris-HCl pH 8.0, 5 mM MgCl2 buffer, before centrifugation again as above. After carefully removing the supernatant, nuclei were used immediately for ATAC-seq.

Detailed methods descriptions for microscopy and time-resolved stomatal conductance measurements can be found in the SI Appendix, Materials and Methods. Detailed protocols for RNA-seq and ATAC-seq library preparation and description of ATAC-seq, RNA-seq, and DAP-seq data analyses can be found in the SI Appendix, Materials and Methods.

### Competing interest statement

The authors declare no competing interests.

## Acknowledgements

Funding for this study was provided by the National Institutes of Health (R01GM060396 to J.I.S and F32GM137544 to C.A.S). CO_2_ experiments were supported by National Science Foundation grant MCB-1900567 to J.I.S. This study includes data generated at the UC San Diego IGM Genomics Center utilizing an Illumina NovaSeq 6000 that was purchased with funding from a National Institutes of Health SIG grant (#S10 OD026929). Microscopy was supported by the UC San Diego Microscopy Core with funding by grant NINDS P30NS047101. We thank Kazuko Yamaguchi-Shinozaki (University of Tokyo) for sharing *abf1/2/3/4* quadruple mutant seeds, and members of the Schroeder lab for helpful discussions throughout the research.

## Author contributions

C.A.S. and J.I.S. conceived of the study and designed the methodology. C.A.S. performed all experiments and analyzed the data. C.A.S. and J.I.S. wrote and edited the paper.

## Data and materials availability

All materials (plasmids, seed stocks) are available upon request. Raw FASTQ files generated in this study and ATAC-seq peak files in narrowPeak format are available at NCBI’s Gene Expression Omnibus under accession GSE243473. All data are available in the main text and supplementary material.

## Notes

### Competing Interest Statement

The authors have declared no competing interest.

### Summary of Updates

Supplemental files updated. Mislabeling of panel in figure 3 was corrected.

