## Supplemental Appendix for "Distinct guard cell specific remodeling of chromatin accessibility during abscisic acid and CO2 dependent stomatal regulation"

**This PDF file includes:**

Supporting text
Figures S1 to S11
Legends for Datasets S1 to S8
SI References

**Other supporting materials for this manuscript include the following:**

Datasets S1 to S8

### Supporting Information Text

### Extended Materials and Methods

#### Microscopy

Plant leaves were mounted on conventional glass microscope slides in water with the abaxial epidermal layer against the coverslip. GFP and chlorophyll fluorescence signals were recorded using either an Eclipse TE2000-U (Nikon, Tokyo, Japan) spinning disc confocal microscope or a Leica SP8 confocal microscope (Leica Microsystems). Constant gain, laser power, and exposure times were used for each experiment when comparing treatments/samples. On the Leica SP8 images were acquired using a 512 x 512 pixel resolution and a scan speed of 400 Hz. Displayed images are representative of at least 4 different leaves harvested from two independent transgenic lines. Images were processed for publication using Fiji software. 3D reconstructions from confocal imaging data were generated using Leica LAS X Software.

#### Time resolved stomatal conductance measurements

For gas exchange experiments, *Col-0* and *abfx4* mutant plants were grown together in the same growth chamber to minimize biological variation. Intact leaves of six-week-old plants were used for gas exchange experiments. Stomatal conductance was measured using a Licor-6400 infrared gas exchange analyzer equipped with a leaf chamber (LI-COR Biosciences). Before beginning measurements, clamped leaves were equilibrated for 45-60 mins at 150 µmol $m^{-2}$ $s^{-1}$ light intensity, ~65% relative humidity, 21ºC, 400 ppm [$CO_2$], and an air flow of 400 µmol $s^{-1}$. Stomatal conductance was recorded every 30 seconds for a total of 30 min. To control for the effect of the diurnal rhythm on stomatal physiology, gas exchange experiments began 2 hours after the beginning of the light cycle and measurements were made alternating between different genotypes. Data are presented as average stomatal conductance +/- standard error of 5 plants per genotype.

#### RNA-seq

For whole leaf RNA-seq, total RNA was extracted from the leaves from 6 plants per sample using the Spectrum total RNA isolation kit. For guard cell enriched RNA-seq, the leaves of 20 plants per sample were blended for 30 sec in ice-cold water, filtered through a 100 µm mesh, and then blended and filtered again as above to isolate epidermal tissue. Thin sections of epidermal tissue were then frozen in liquid nitrogen and ground to a fine powder using a mortar and pestle. Total RNA was then extracted from ground tissue powder using the Qiagen RNeasy mini kit with on-column DNaseI digestion. mRNA-seq libraries were prepared and sequenced on a NovaSeq 6000 (PE100) at the UCSD Institute for Genomic Medicine.

Following sequencing, adapter trimming and quality control checks were performed using Trim Galore version 0.6.5 (Cutadapt and FastQC). Reads were aligned to the *Arabidopsis thaliana* genome (TAIR10) using the Rsubread (version 2.14.0) *align* function (1). Low quality alignments (mapping quality < 2) were removed using the Samtools *view* function (version 1.17) (2). RNA-seq reads per transcript were counted to generate count matrices using the Rsubread *featureCounts* function. Differential expression analysis was performed using DESeq2 (version 1.26.0) (3). Transcripts were defined as significantly differentially expressed using the following thresholds: P adjusted value < 0.001 and fold change > 2. To visualize data using the Integrated Genomics Viewer (IGV) browser, bigwig coverage tracks were generated using the deepTools (version 3.5.0) *bamCoverage* function (bin size of 1 bp and normalization by Reads Per Kilobase per Million mapped reads - RPKM) (4). Gene ontology (GO) term enrichment analyses were performed with the Panther system and by applying a binomial test with p-values adjusted for multiple testing by a Bonferroni correction (5).

#### ATAC-seq

ATAC-seq was performed as described in (6). Sorted nuclei were resuspended in 40 µl of tagmentation reaction mixture (20 µl 2xTD buffer, 0.4 µl 10% Tween, 0.4 µl 1% digitonin, 13.2 µl 1xPBS, X µl H2O) supplemented with TDE1 enzyme (Illumina) at a ratio of 2.0 µl TDE1:50,000 nuclei. Reactions were incubated at 37C for 30 mins. After incubation, tagmented DNA fragments were purified using the Qiagen min-Elute DNA kit. ATAC-seq libraries were amplified with dual indexed Nextera primers using the NEBNext 2xHiFi PCR Master Mix (New England Biolabs) for 5 initial cycles. The number of additional amplification cycles was determined empirically by performing qPCR on 10% of the amplified library. Amplified libraries were then dual size selected using a 0.6x-1.2X ratio of SPRIselect magnetic beads (Beckman Coulter) and library quality was assessed by running samples on a HSD1000 ScreenTape on a Tapestation system (Agilent). ATAC-seq libraries were pooled at equimolar ratios and sequenced on a NovaSeq 6000 (PE150 mode) by Novogene.

To control for differences in data analysis between studies, previously published Mesophyll cell ATAC-seq raw FASTQ files were downloaded from SRA (SRP113667) and reanalyzed. Raw ATAC-seq sequence files were subjected to adapter trimming and quality control checks using Trim Galore version 0.6.5 (Cutadapt and FastQC). Throughout the computational analysis parallel processing was performed using GNU Parallel (7). Processed reads were aligned to the *Arabidopsis thaliana* genome (TAIR10) using the Rsubread *align* function (version 2.14.0). Low quality alignments (mapping quality < 2) and alignments to the mitochondrial and chloroplast genomes were removed using the Samtools *view* function (version 1.17). Alignments in a region of chromosome 2 (3239001 – 3510171) with abnormally high ATAC-seq signal in all samples were removed prior to further processing using BEDtools (version 2.30) (8). Reads derived from PCR duplicates were marked for exclusion using the Picard tools function *Markduplicates* (version 1.141). ATAC-seq peaks were called across all biological replicated using Genrich (9) with the following parameters: -r -d -j -p 0.01 -a 200. Peaks were then merged across all samples using the BEDtools *merge* function to generate a master list of peaks. ATAC-seq alignments were then assigned to peaks and counted using the Rsubread *featureCounts* function and the output from this function was used to calculate Fraction of Reads in Peaks scores (FRiPs) for each library. Differential chromatin accessibility analysis was performed using DESeq2 (version 1.26.0). Regions were defined as significantly differentially accessible using the following thresholds: P adjusted value < 0.001 and fold change > 1.5 or 2 as indicated in associated figure legends. To visualize data using the Integrated Genome Browser (IGV), bigwig coverage tracks were generated on merged replicate bam files using the deepTools (version 3.5.0) *bamCoverage* function (bin size of 1 bp and normalization by RPKM). Homer software (version 2.1.2) (10) was used to annotate ACRs and to assign ACRs to the nearest downstream transcript (within -2.5 kb and +0.5 kb). Gene ontology (GO) term enrichment analyses were performed with the Panther system, and by applying a binomial test with p-values adjusted for multiple testing by a Bonferroni correction. Heatmaps and metaplots of were generated with DeepTools. Homer software was used for de novo motif discovery among differentially accessible ACRs and to match discovered motifs to previously known motifs from published databases. We used ChIPpeakAnno software (Version 3.24) (11) to assign ACRs to annotated genomic features and to test for genomic overlap among different sets of peaks.

**DAP-seq Analysis**
To control for differences in data analysis between studies, previously published DAP-seq datasets GSE60143 (12) and PRJNA682697 (13) were reanalyzed. Raw FASTQ files were downloaded from SRA and processed to remove adapters using Trim Galore. Reads were aligned to the *Arabidopsis thaliana* genome using the Rsubread *align* function and low-quality alignments (mapping quality < 2) were removed using the Samtools *view* function. DAP-seq peaks were called using the MACS2 *callpeak* function with the following parameters: –keep-dup 1 --gsize 1.2e8 (version 2.2.7). For ABF1/2/3/4 DAP-seq datasets, peaks were merged across four replicate experiments. Bigwig files were generated on merged DAP-seq bam files using the deepTools (version 3.5.0) *bamCoverage* function (bin size of 1 bp and normalized via RPKM). DeepTools software was used to generate heatmaps and Metaprofile plots centered over ABA-regulated ACRs.

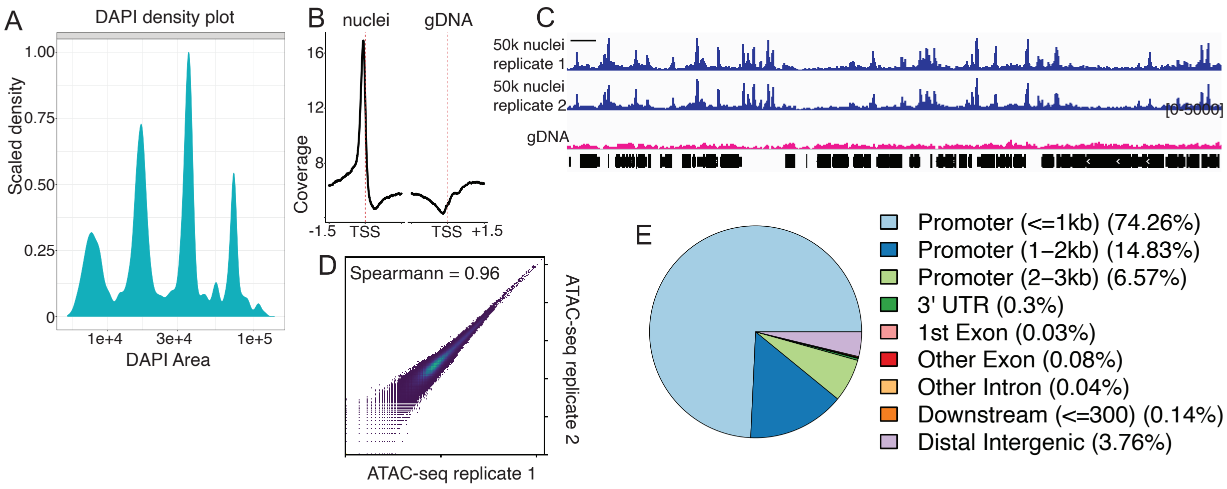


**Fig. S1. FANS-ATAC-seq on seedling nuclei reproducibly assays chromatin accessibility in regulatory DNA. A)** Density plot of DAPI fluorescence shows the distribution of ploidy among FACS-sorted nuclei from seedlings. **B)** Open chromatin reads (<100 bp) of a seedling ATAC-seq library shows strong enrichment upstream of transcription start sites (TSSs). **C)** Genome browser snapshot of a region of chromosome 1 showing ATACseq signal derived from either 50,000 seedling nuclei (two replicates shown in blue) or from purified genomic DNA (in pink). **D)** Scatterplot of ATAC-seq coverage showing correlation between two ATAC-seq replicates. **E)** Genomic distribution of seedling nuclei ATAC-seq peaks among annotated features.

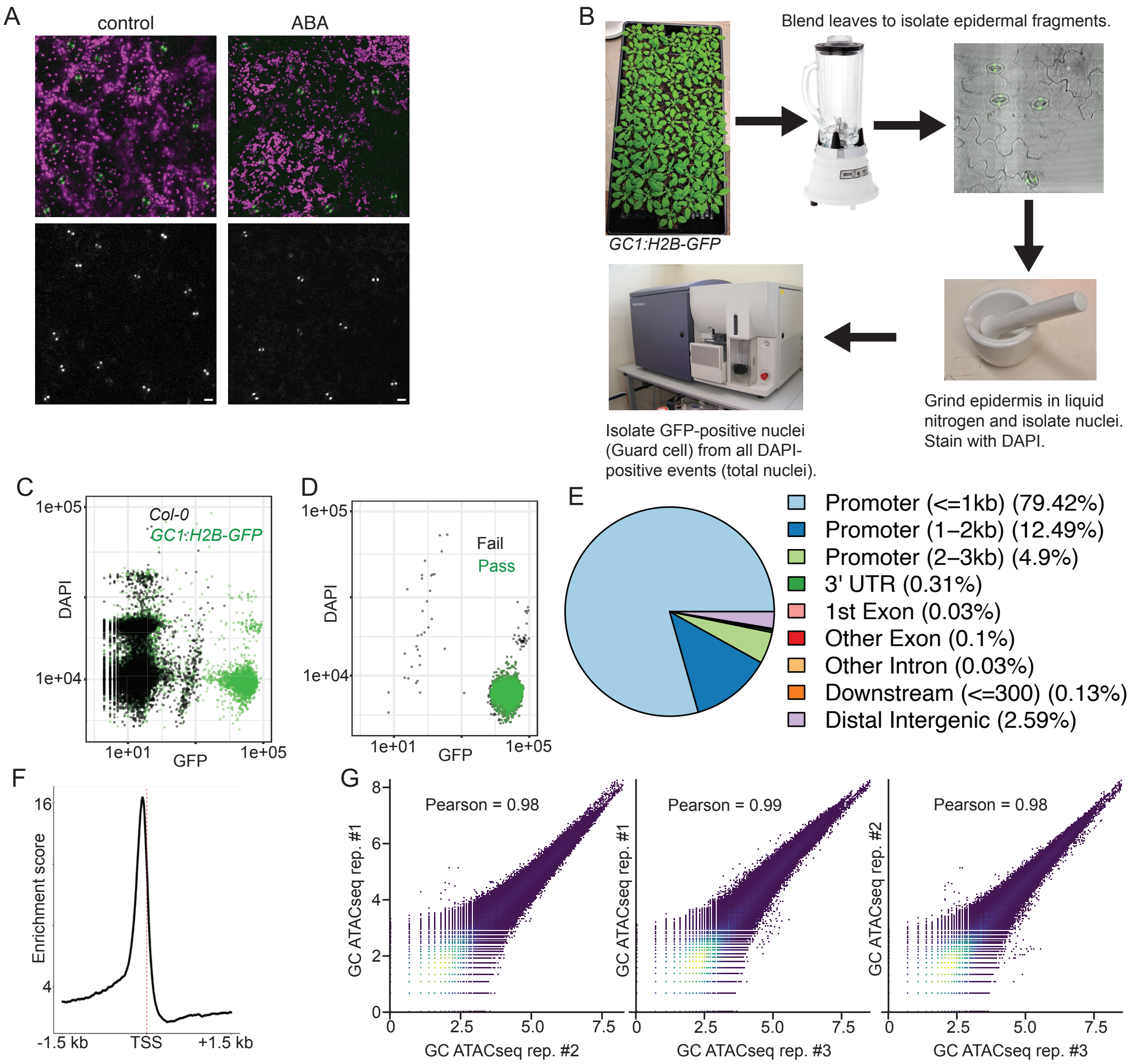


**Fig. S2. Guard cell FANS-ATAC-seq. A)** Micrographs from confocal imaging of 5-week-old *GC1p:H2B-GFP* leaves treated with either control or ABA for 4hr. Images on top show merged GFP (green) and chlorophyll fluorescence (magenta). Bottom images show GFP signal only. **B)** Diagram of the protocol used to isolate guard cell nuclei from *Arabidopsis* plants. The leaves of roughly 40 6-week-old *GC1:H2b-GFP* plants were blended in ice cold water. The resulting blendate was filtered through a 100 µm mesh sheet to isolate epidermal tissue. A representative fluorescence micrograph shows intact guard cells with nuclei labeled by H2b-GFP. Epidermal tissue was flash frozen in LN2 and then ground to a fine powder using a mortar and pestle. After extracting nuclei from ground tissue, GFP positive nuclei were purified using FACS. **C)** DAPI vs GFP intensity plot resulting from FACS analysis of nuclei from either *GC1:H2b-GFP* plants (in green) or *Col-0* plants (in black). **D)** Sorted guard cell nuclei were re-run through the same FACS protocol to evaluate the purity of the isolated population. The DAPI vs GFP plot shows nuclei passing (green) and failing (black) the sorting criteria. Out of 10,000 sorted nuclei, 9821 passed (purity of 98%). **E)** Pie chart displaying the distribution of guard cell ATAC-seq peaks among annotated genomic features. **F)** TSS enrichment plot of open chromatin reads (<100 bp) from a Guard Cell ATAC-seq library shows strong enrichment upstream of transcription start sites (TSSs). **G)** Scatterplots showing the correlation among biological replicates of guard cell ATAC-seq libraries.

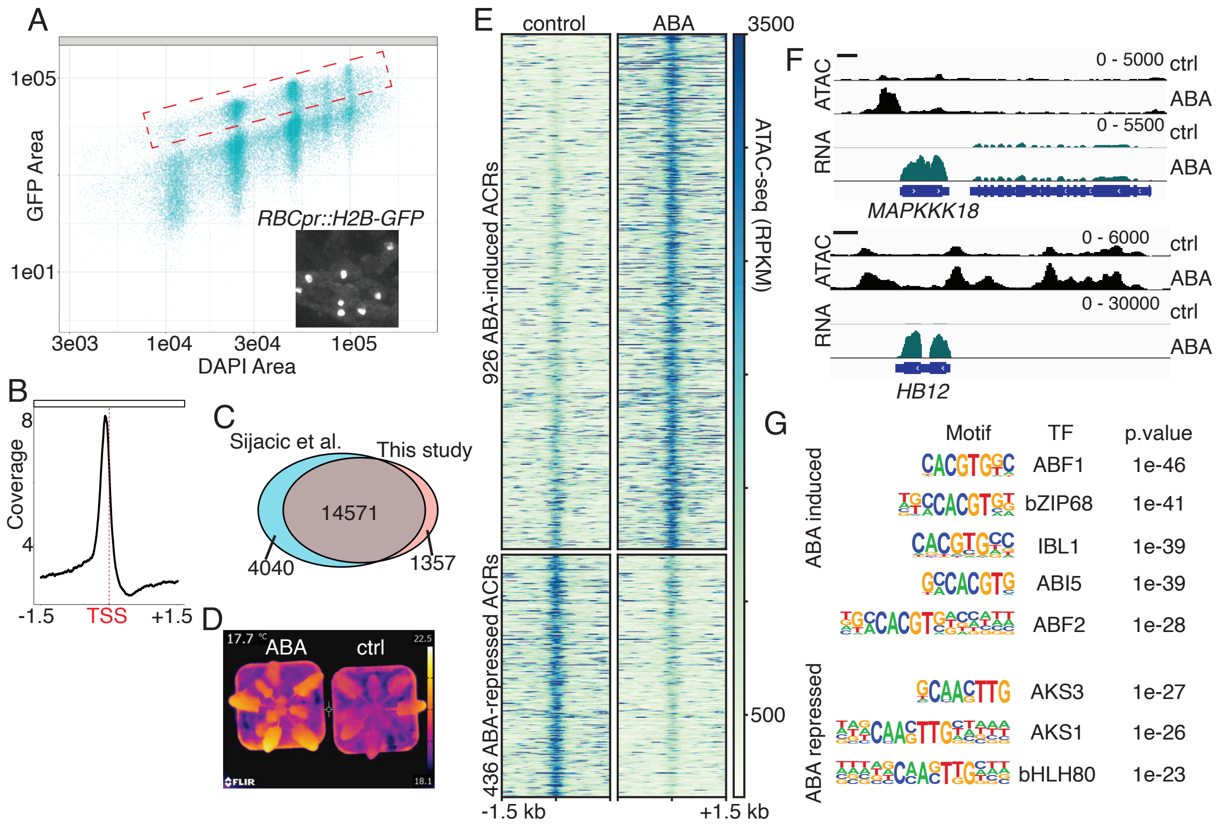


**Fig. S3. ABA induced chromatin remodeling in mesophyll cell nuclei. A)** GFP vs DAPI intensity plot showing the results of FACS analysis on nuclei from *RBCp::H2b-GFP* plants. The location of GFP positive mesophyll nuclei sorted for ATAC-seq is indicated dashed box. Inset shows a confocal image showing the expression of *RBCp::H2b-GFP* in mesophyll cell nuclei. **B)** ATAC-seq coverage plot showing strong enrichment of mesophyll ATAC-seq reads upstream of Transcription Start Sites (TSSs). **C)** Overlap of mesophyll ATAC-seq peaks from (14) with those from this study. **D)** Representative IR image showing increase in leaf surface temperature 3hrs after spraying *RBC:H2b-GFP* plants with ABA. **E)** Heatmap of ATAC-seq signal in mesophyll nuclei at regions showing ABA-regulated chromatin accessibility. ABA significantly increased accessibility at 926 regions and decreased accessibility at 436 regions (FDR < 0.001 and FC > 1.5). **F)** Genome browser snapshots at representative genes (*MAPKKK18* and HB12) showing ABA-regulated chromatin accessibility in upstream sequences. **G)** Selection of transcription factor binding motifs enriched in ABA-induced and ABA-repressed ATAC-seq regions in mesophyll nuclei.

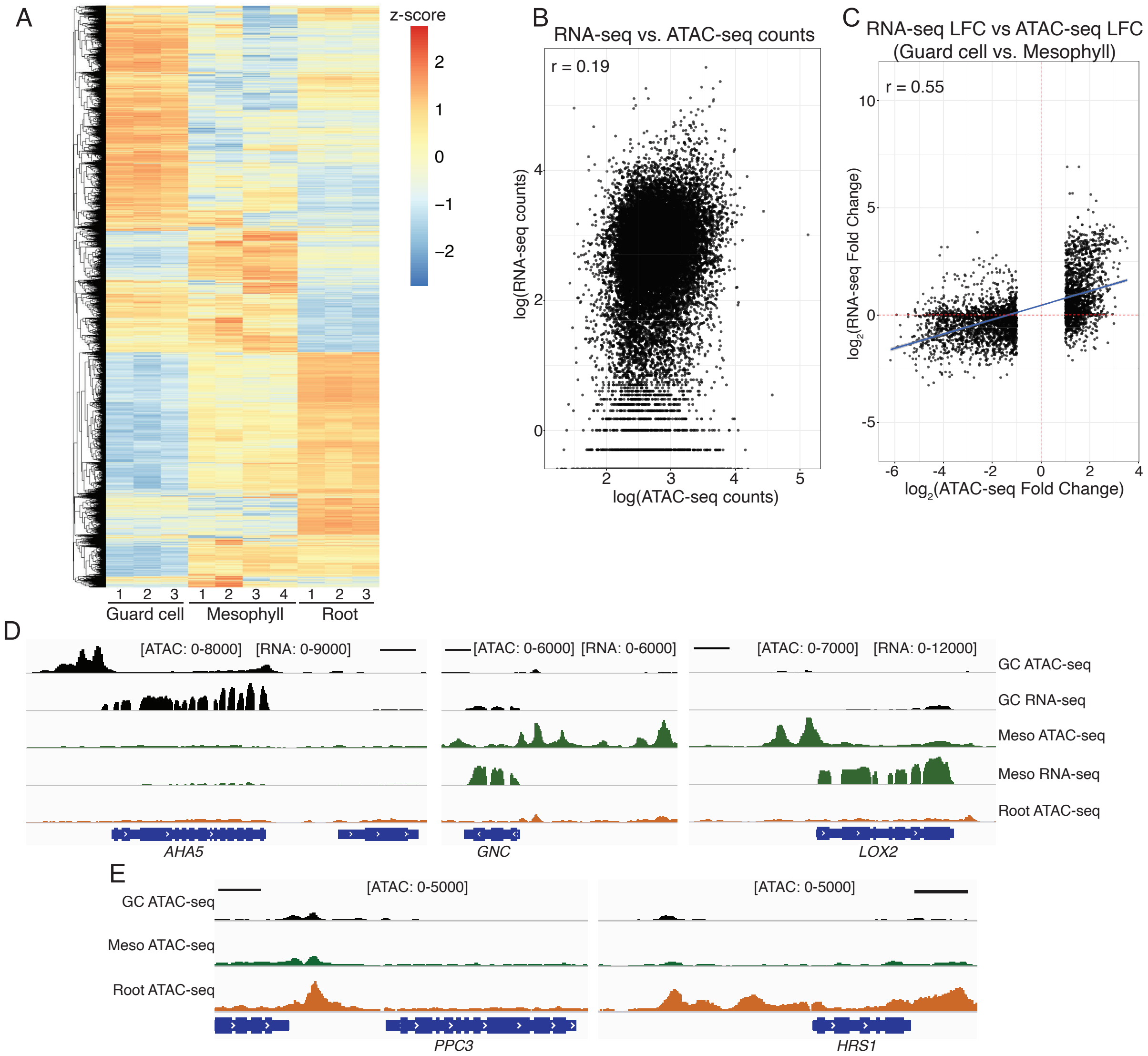


**Fig. S4. Patterns of chromatin accessibility in guard cells, mesophyll cells, and roots. A)** Clustered heat map of normalized ATAC-seq signal plotted over all ACRs identified across samples (30,985 peaks). The indicated samples are split by biological replicate, and ATAC-seq signal is normalized by z-score across rows. **B)** Scatterplot of upstream chromatin accessibility (ACRs between -2.5 kb and +0.5 kb of a transcription start site) and downstream gene transcript levels. ATAC-seq and RNA-seq values are plotted as log(counts). The Pearson's correlation coefficient derived from this comparison is indicated on the plot. **C)** Scatter plot showing the relationship between fold-change in ATAC-seq signal at cell-type regulated ACRs (Guard Cell vs. Mesophyll) and fold-changes in downstream transcript levels. ACRs residing within -2.5 kb and +0.5 kb of a transcription start site were assigned to nearest gene. A linear trend line (blue) with 95% confidence interval (in gray) is shown. The Pearson's correlation coefficient derived from this comparison is indicated on the plot. **D)** IGV browser displays of the indicated ATAC-seq and RNA-seq data highlighting examples of cell-type specific accessible chromatin, scale bars represent 1 kilobase. Guard cell specific open chromatin upstream of the plasma membrane H+-ATPase gene *AHA5.* Mesophyll-specific open chromatin upstream of the GATA transcription factor gene *GNC*. Mesophyll-specific open chromatin upstream of the lipoxygenase gene *LOX2*. **E)** IGV browser displays of the indicated ATAC-seq data highlighting examples of root specific open chromatin upstream of the phosphoenolpyruvate carboxylase gene *PPC3* and upstream of the GARP-family transcription factor gene *HRS1.* Scale bars indicate 1 kilobase.

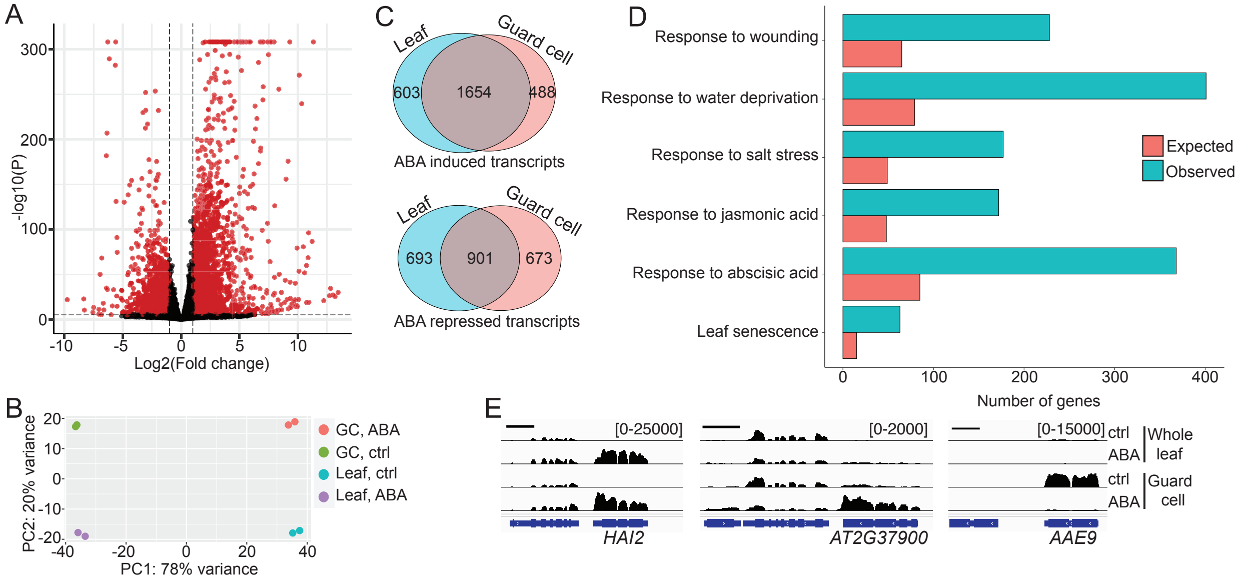


**Fig. S5. RNA-seq identifies ABA-regulated transcripts in guard cells and whole leaf tissue.** **A)** Volcano plot of differential whole leaf RNA-seq analysis (FDR < 0.001, FC > 2) comparing control and ABA-treated samples. Differentially expressed transcripts are colored in red (2257 upregulated and 1594 downregulated). **B)** Principal component analysis (PCA) of RNA-seq results showing samples clustering by treatment (ABA vs control) and cell-type (guard cell vs leaf). **C)** Overlap of transcripts induced or repressed by ABA between whole leaves and guard cells. **D)** Results of a Gene Ontology (GO) analysis performed on ABA-induced transcripts in guard cells. **E)** Genome browser snapshots showing RNA-seq signal from the indicated samples at the representative ABA-regulated genes *HAI2, AT2G37900,* and *AAE9*.

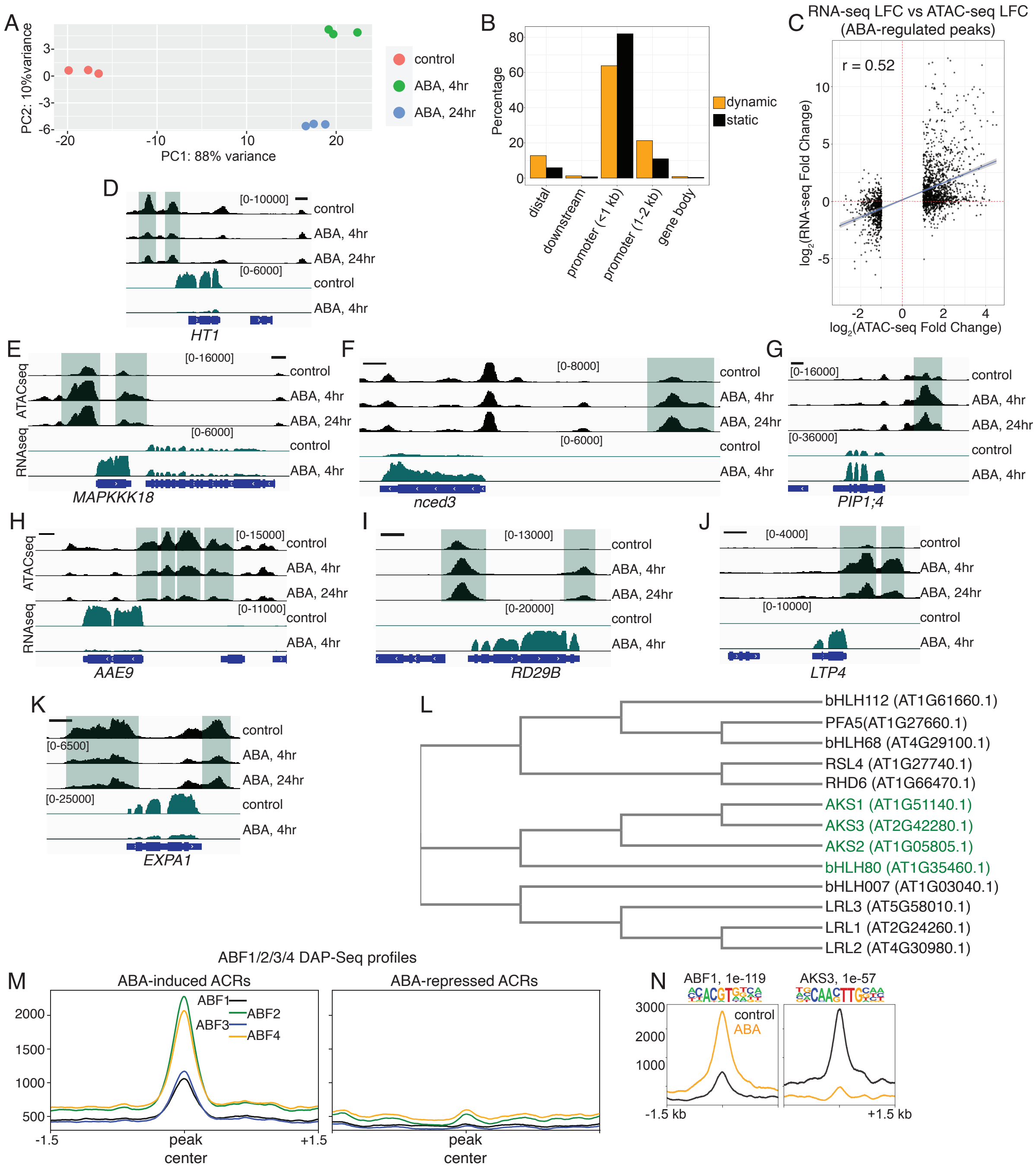


**Fig. S6. ABA triggered chromatin remodeling in guard cells. A)** PCA plot of guard cell ATAC-seq samples from ABA treatment experiments. **B)** Genomic distribution among annotated features of guard cell ATAC-seq peaks separated into regions that are static (black) and those that are ABA-regulated (orange). **C)** Scatter plot showing the relationship between fold-change in ATAC-seq signal at ABA-regulated ACRs in guard cells and fold-change in downstream transcript levels in guard cells. ACRs residing within -2.5 kb and +0.5 kb of a transcription start site were assigned to nearest gene. A linear trend line (blue) with 95% confidence interval (in gray) is shown. The Pearson's correlation coefficient derived from this comparison is indicated on the plot. **D-K)** IGV images (scale bars indicate 1 kb) showing additional examples of ABA-induced changes to chromatin accessibility upstream of **D)** *HT1* which encodes a kinase controlling low $CO_2$ induced stomatal opening, **E)** the ABA-activated kinase *MAPKKK18*, **F)** ABA biosynthesis gene *NCED3*, **G)** the wax biosynthesis gene *AAE9*, **H)** *RD29B*, **I)** the aquaporin *PIP1;4*, **J)** *LTP4*, and **K)** the cell wall expansin gene *EXPA1*. **L)** Cladogram generated from multiple sequence alignment (ClustalW) of related *Arabidopsis* bHLH transcription factors. TFs with motifs enriched in ABA-repressed ACRs are highlighted in green. **M)** Metaprofile plots of DAP-seq signal (RPKM-normalized) for ABF1/2/3/4 centered over ABA-induced or ABA-repressed ACRs. **N)** De novo motif discovery uncovered motifs similar to ABF1 and AKS3 in ABA-activated and ABA-repressed ACRs, respectively. The p-value associated with this enrichment is shown. Below the uncovered motifs are metaplots of ATAC-seq signal (RPKM-normalized) from either control or ABA-treated guard cells centered over peaks containing these motifs.

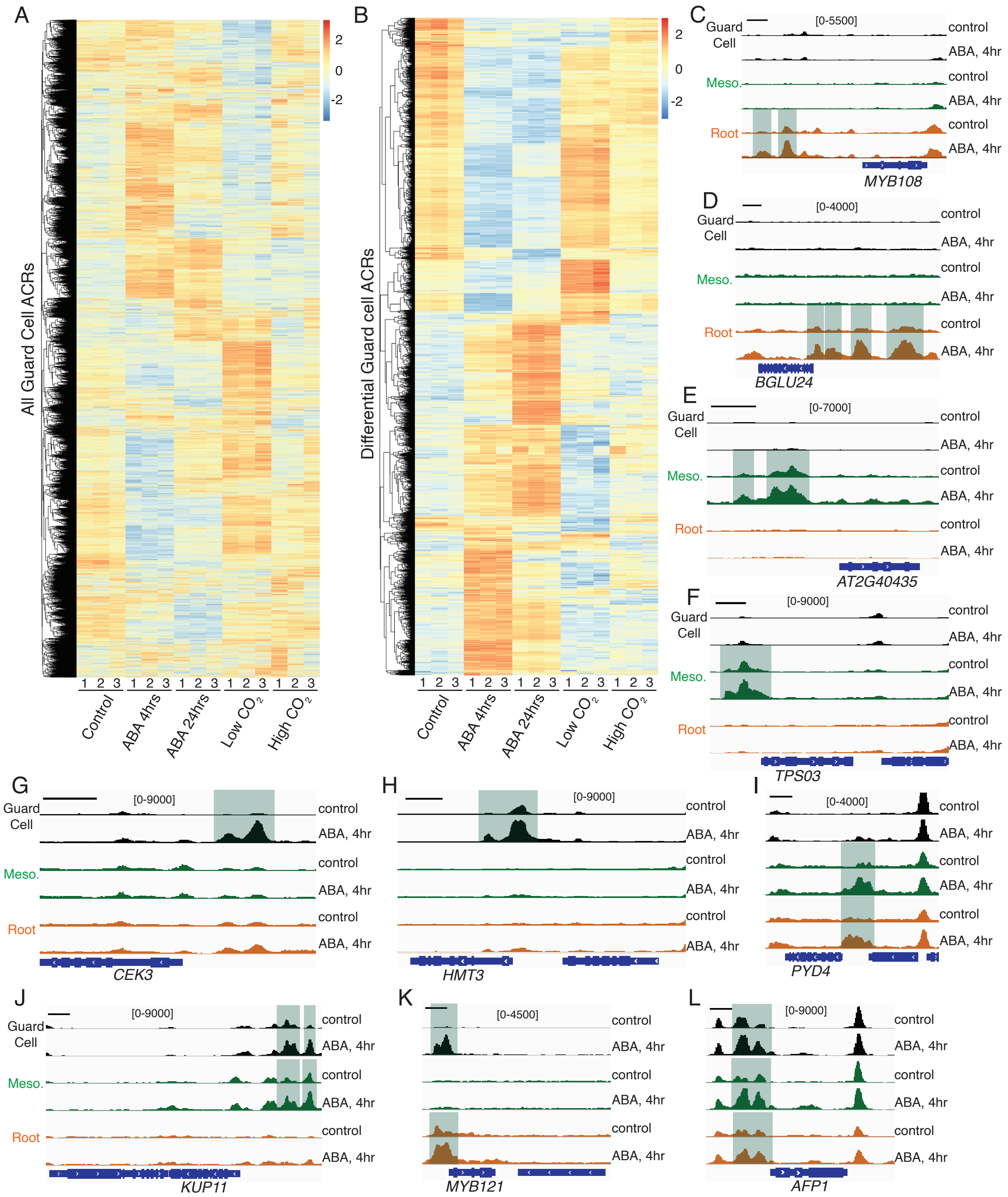


**Fig. S7. Chromatin dynamics in guard cells and examples of cell-type specificity. A)** Clustered heat map of normalized ATAC-seq signal plotted over all ACRs (22680 peaks) identified in guard cells. The indicated samples are split by biological replicate, and ATAC-seq signal is normalized by z-score across rows. **B)** Clustered heat map of normalized guard cell ATAC-seq signal plotted over differential ACRs across all treatments (4473 peaks). The indicated samples are split by biological replicate, and ATAC-seq signal is normalized by z-score across rows. **C-L)** Genome browser images showing examples of cell/tissue-type specific ABA chromatin dynamics. Scale bars represent 1 kilobase and the relative positions of differential ACRs are indicated by green highlighting. Root specific changes to chromatin accessibility upstream of **C)**

the MYB transcription factor gene *MYB108* and **D)** the beta glucosidase gene *BGLU24.* Mesophyll-specific ABA-induced chromatin dynamics upstream of **E)** the SCREAM-like gene *AT2G40435* and **F)** the terpene synthase gene *TPS03*. Guard cell specific ABA-induced chromatin dynamics upstream of **G)** the choline synthase gene *CEK3* and **H)** the homocysteine S-methyltransferase gene *HMT3.* Examples of ABA-induced chromatin dynamics common to multiple cell/tissue-types upstream of **I)** the gene *PYD4,* **J)** the potassium transporter gene *KUP11,* **K)** the MYB-transcription factor gene *MYB121,* and **L)** the ABI5 binding protein gene *AFP1*.

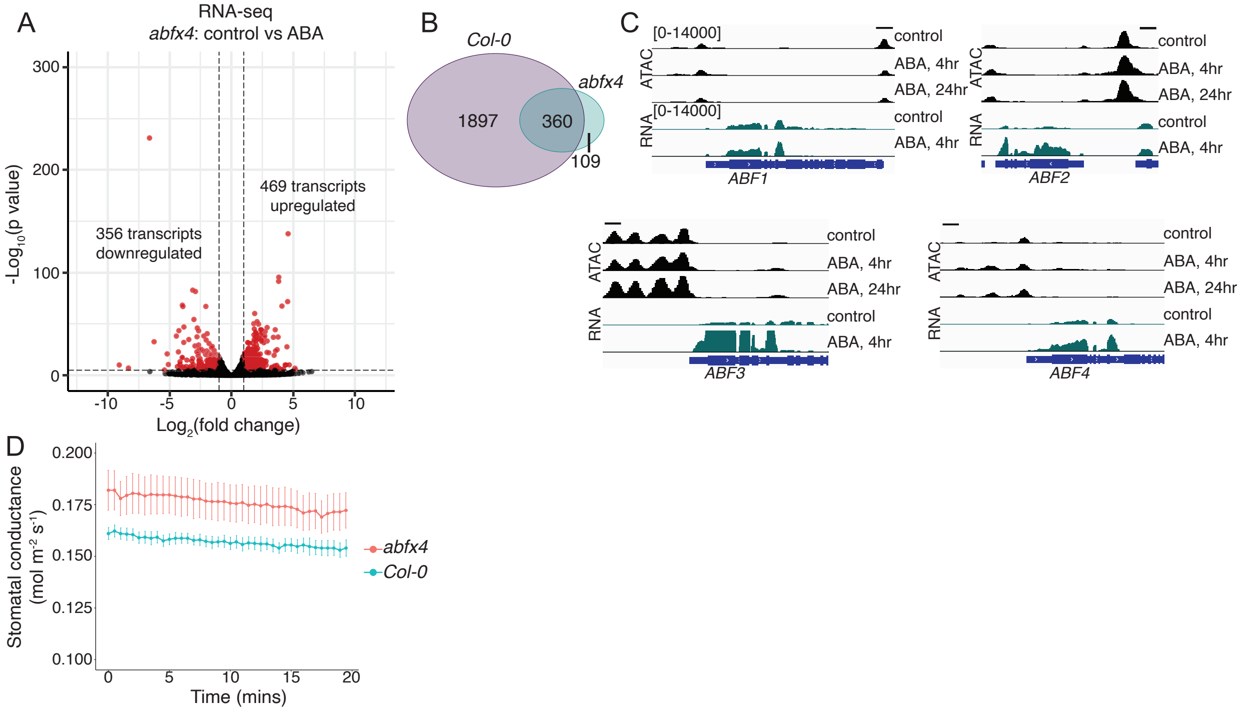


**Fig S8. The ABRE binding proteins ABF1-4 are required for the bulk of ABA-induced transcription in mature leaf tissue. A)** Volcano plot of RNA-seq results from ABA or control treated 5-week-old *abfx4* mutant leaf tissue. ABA significantly upregulated 469 transcripts and downregulated 356 transcripts (FDR < 0.001 and FC > 2). **B)** Overlap of ABA-induced transcripts in *Col-0* leaves vs *abfx4* mutant leaves shows that ABFs are required for the upregulation of 2030 transcripts. **C)** Genome browser snapshots showing guard cell ATAC-seq signal and RNA-seq signal at four genes encoding ABF proteins. Chromatin accessibility upstream of *ABF1-4* was not significantly increased by ABA in guard cells. **D)** Gas exchange (Licor-6400) comparing steady state stomatal conductance of well-watered *Col-0* and *abfx4* plants (n = 5 per genotype).

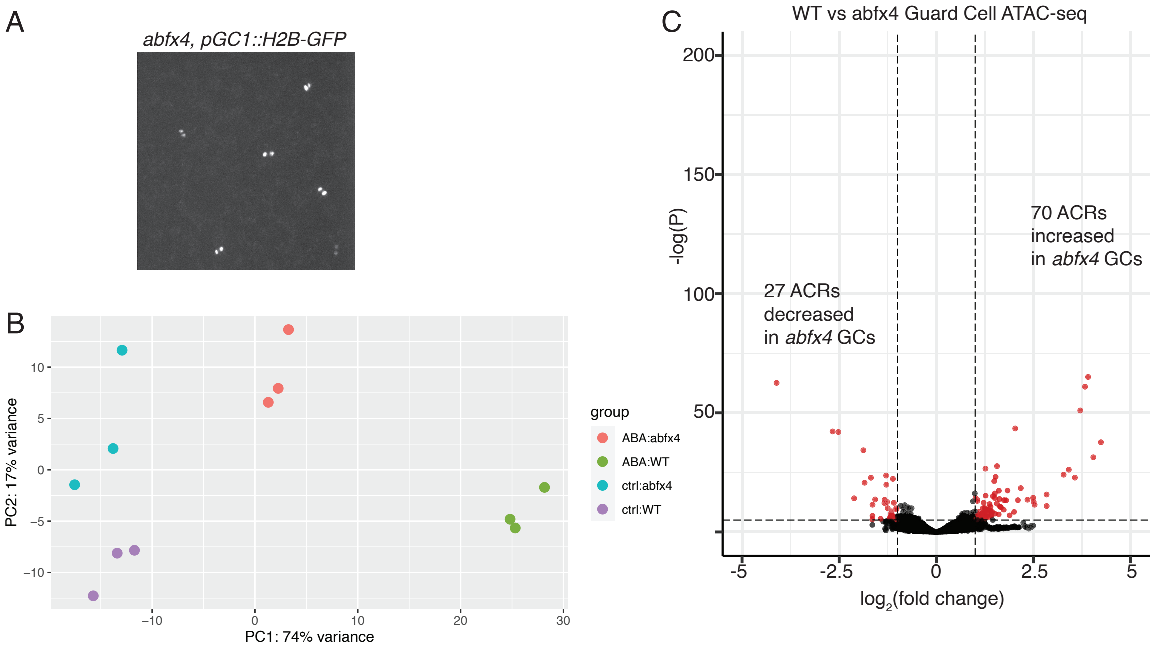


**Fig S9. Chromatin accessibility in *abfx4* mutant guard cells without ABA treatment. A)** Confocal image showing guard cell specific expression of H2B-GFP in *abfx4* mutant leaf. **B)** Principal component analysis (PCA) of Guard cell ATAC-seq results showing samples clustering by treatment (ABA vs control) and genotype (*Col-0* vs *abfx4*). **C)** Volcano plot summarizing differential ATAC-seq analysis (FDR < 0.001 and FC > 2.0) comparing chromatin accessibility in control treated *WT* vs *abfx4* mutant guard cells.

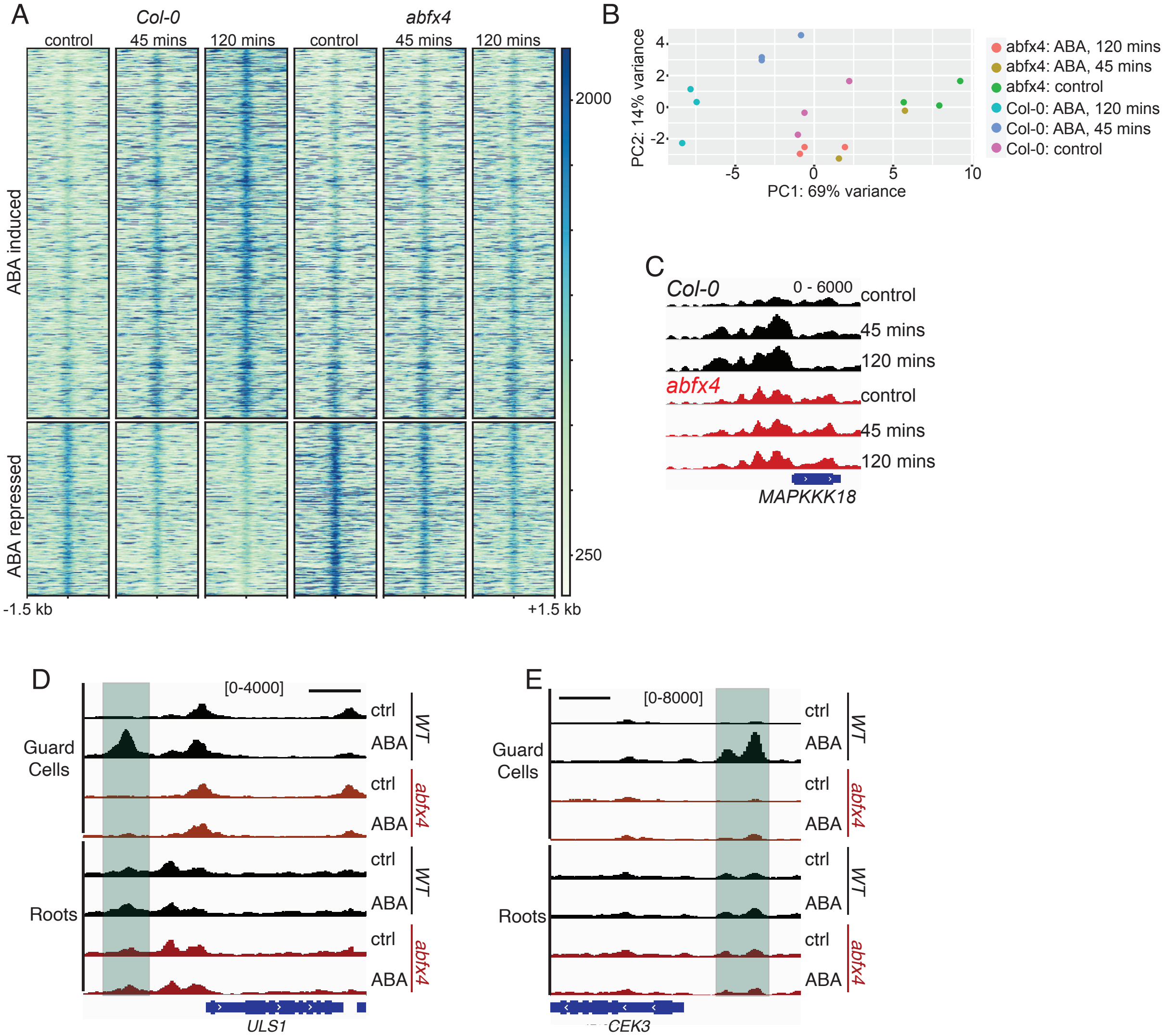


**Fig S10. ABFs are required for ABA-induced chromatin opening in roots. A)** Heatmap of ATAC-seq signal at regions showing ABA-regulated chromatin accessibility over the indicated ABA treatment time course in either *WT* or *abfx4* mutant root nuclei. **B)** Principal component analysis of root ATAC-seq libraries showing separation of samples by both genotype and ABA treatment duration. **C)** Genome browser snapshot showing chromatin accessibility in *WT* or *abfx4* mutant root nuclei upstream of the ABA-induced gene *MAPKKK18* over the indicated ABA treatment time series. **D, E)** Genome browser snapshots showing guard cell specific and ABF-dependent ABA-induced ACRs upstream of the genes *CEK1 and ULS1.* Scale bars indicate 1 kb.

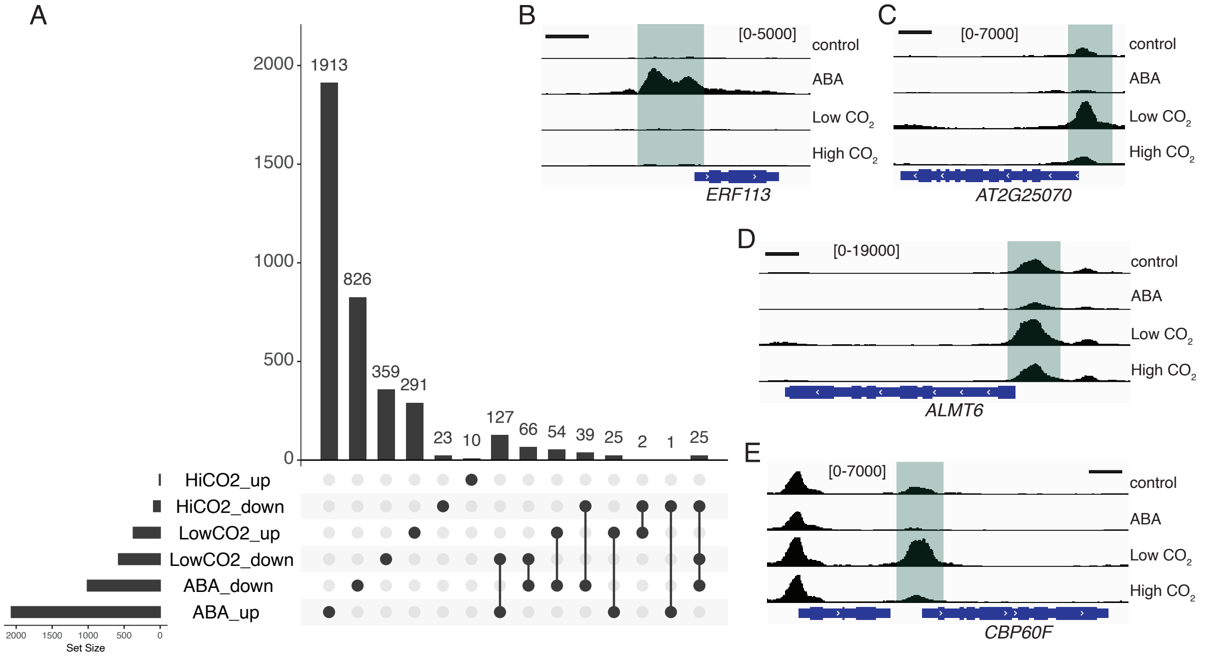


**Fig S11. Distinct regulation of chromatin by ABA and $CO_2$ in guard cells. A)** UpSet plot showing the set relationships between ABA, low $CO_2$, and high $CO_2$ regulated ATAC-seq peaks. The y-axis of the bar-plot shows the number of peaks found in the set intersections indicated along the x-axis. The sets being comparing are indicated by lines connecting filled in circles. **B-E)** Genome browser images (scale bars indicate 1 kb) of ATAC-seq signal (RPKM-normalized) showing ABA and $CO_2$-regulated chromatin accessibility upstream of **B)** *ERF113*, **C)** *AT2G25070*, **D)** *ALMT6*, and **E)** *CBP60F*.

### Supplementary Datasets

**Dataset S1**. Summary of sequencing libraries. Excel file containing 1 sheet.

**Dataset S2**. Annotated lists of all ATAC-seq peaks called in different cell/tissue-types. Annotations include genomic coordinates, distance to nearest transcription start site, and nearest gene name and description. Excel file containing 10 sheets: **1)** Annotated list of all ACRs called in root nuclei ATAC-seq libraries. **2)** Annotated lists of all ACRs called in Guard Cell nuclei ATAC-seq libraries. **3)** Annotated list of all ACRs called in Mesophyll cell nuclei ATAC-seq libraries. **4)** Annotated list of all ACRs called in Mesophyll cell ATAC-seq libraries from (14). **5)** Annotated list of ACRs enriched in Root nuclei. **6)** Annotated list of ACRs enriched in Mesophyll cell nuclei. **7)** Annotated list of ACRs enriched in Guard Cell nuclei. **8)** Master list of merged ATAC-seq peaks from called across all tissues/cell-types. **9)** Master list of merged ATAC-seq peaks called in root ATAC-seq samples across all treatments and genotypes. **10)** Master list of merged ATAC-seq peaks called in guard cell ATAC-seq samples across all treatments and genotypes.

**Dataset S3**. Annotated lists of all differentially accessible ACRs. Annotations include genomic coordinates, distance to nearest transcription start site, and nearest gene name and description. Datasets also contain results of differential chromatin accessibility analysis including log2(Fold-change) and adjusted p-values. Excel file containing 13 sheets: **1)** Annotated differentially accessible ACRs in whole seedings (control vs. 4 hours ABA). **2)** Annotated differentially accessible ACRs in roots (control vs. 45 minutes ABA). **3)** Annotated differentially accessible ACRs in roots (control vs 2 hours ABA). **4)** Annotated differentially accessible ACRs in roots (control vs 4 hours ABA). **5)** Annotated differentially accessible ACRs in mesophyll nuclei (control vs 4 hours after ABA treatment). **6)** Annotated differentially accessible ACRs in guard cell nuclei (control vs 4 hours after ABA treatment). **7)** Annotated differentially accessible ACRs in guard cell nuclei (control vs 24 hours after ABA treatment). **8)** Annotated differentially accessible ACRs in guard cell nuclei (no treatment, WT vs. *abfx4* mutant). **9)** Annotated differentially accessible ACRs in *abfx4* mutant guard cell nuclei (control vs. 4 hours after ABA treatment). **10)** Annotated differentially accessible ACRs in *abfx4* mutant root nuclei (control vs. 45 minutes ABA). **11)** Annotated differentially accessible ACRs in *abfx4* mutant root nuclei (control vs 2 hours ABA). **12)** Annotated differentially accessible ACRs in guard cell nuclei (control vs. 100 ppm $CO_2$). **13)** Annotated differentially accessible ACRs in guard cell nuclei (control vs 1000 ppm $CO_2$).

**Dataset S4**. Table containing annotated high confidence guard cell specific ACRs ranked by downstream gene transcript level in guard cells. Excel file containing 1 sheet.

**Dataset S5**. ABA-regulated transcripts in whole leaves and guard cells. Excel file containing 2 sheets: **1)** Differentially expressed genes in whole leaves (control vs. 4 hours after ABA treatment). **2)** Differentially expressed genes in guard cells (control vs 4 hours after ABA treatment).

**Dataset S6**. Table containing results of Gene Ontology (GO) term analyses of genes downstream of ABA-regulated ACRs. Excel file containing 6 sheets: **1)** GO terms enriched among genes downstream of ABA (4 hr)-induced ACRs in roots. **2)** GO terms enriched among genes downstream of ABA (4 hr)-repressed ACRs in roots. **3)** GO terms enriched among genes downstream of ABA (4 hr)-induced ACRs in guard cells. **4)** GO terms enriched among genes downstream of ABA (4hr)-repressed ACRs in guard cells. **5)** GO terms enriched among genes downstream of ABA (24 hr)-induced ACRs in guard cells. **6)** GO terms enriched among genes downstream of ABA (24 hr)-repressed ACRs in guard cells.

**Dataset S7**. Table containing Top 10 transcription factor binding motifs identified among different sets of differentially accessible ATAC-seq peaks.

**Dataset S8**. Table of primer sequences. Excel file containing 1 sheet.
